# Mouse strain and network-level activity differences underlie social decision-making

**DOI:** 10.64898/2026.03.06.710206

**Authors:** Elizabeth Illescas-Huerta, Andrea Villamizar, Meghan Cum, Nancy Padilla-Coreano

## Abstract

Adaptive social behavior requires balancing self-interest with the welfare of others, a core axiom of social decision-making that determines whether actions are selfish or prosocial. Although the medial prefrontal cortex (mPFC) has been implicated in prosocial behavior, the broader cortical-subcortical networks that arbitrate between selfish and prosocial actions remain poorly understood. Moreover, most studies of social decision-making (at both the single-region and circuit levels) have focused on inbred C57BL/6 mice, leaving unclear whether similar neural mechanisms operate across genetically diverse populations. Here, we combined a social decision-making task with c-Fos mapping to examine activity across distributed cortical and subcortical regions in inbred C57BL/6 and outbred CD1 male mice during prosocial and selfish choices. We found that CD1 mice exhibited a stronger bias toward selfish behavior, whereas C57BL/6 mice were more prosocial. This behavioral divergence was associated with elevated c-Fos activity in the mPFC and nucleus accumbens core (NAcC) in CD1 mice compared with C57BL/6 mice, and mPFC activity positively correlated with selfish choice bias. At the network level, social decision-making selectively recruited coordinated activity among the distinct mPFC subregions, ventral tegmental area (VTA), and NAcC. Importantly, prosocial and selfish individuals recruited distinct prefrontal-subcortical network configurations during social decision-making. Together, these findings identify distributed cortical-subcortical network dynamics underlying social choice bias and reveal strain-dependent differences in the neural architecture supporting prosocial and selfish behavior.

**Significant statement:** Social decisions require weighing personal benefit against the welfare of others, yet the neural circuits that bias individuals toward selfish versus prosocial choices remain poorly understood. Here, we show that two mouse strains with opposing social preferences recruit distinct cortical and subcortical network configurations during social choice, despite performing the same task. Rather than reflecting differences in single brain regions, social decision-making engaged coordinated activity across a prefrontal–striatal–midbrain circuit, with prosocial and selfish individuals recruiting different versions of this network. These findings reveal that social choice bias is encoded at the level of distributed circuit organization and that genetic background shapes how the brain implements social decisions.

## INTRODUCTION

In social contexts, adaptive behavior depends on the ability to integrate self-interest with the interests of others, a central component of social decision-making (Rilling and Sanfey, 2011). Within this framework, social decisions can bias behavior toward prosocial choices—defined as intentional actions that confer benefits to others—or selfish actions that prioritize individual gain. Prosocial decisions are conserved across species and play a critical role in maintaining social cohesion and supporting group survival (Rault 2019; Wu an d Hong 2022). In rodents, prosocial behaviors such as cooperation, harm avoidance, helping, and reward provision have been widely used to investigate the neural basis of social decision-making (Keysers et al. 2022; Michael J.M. Gachomba et al. 2024). By contrast, the neural mechanisms that bias decisions toward selfish outcomes, including the brain’s mechanisms for arbitrating between prosocial and selfish actions, remain largely unknown.

The medial prefrontal cortex (mPFC) has emerged as a central hub for social decision-making (Gangopadhyay et al. 2021), positioning it as a key region for arbitrating between prosocial and selfish actions. Within the mPFC, distinct subregions support different expressions of prosocial behavior in mice. Activity in the anterior cingulate cortex (ACC) is necessary for harm avoidance toward conspecifics (Song et al. 2023), and plays a role in empathy-like behavior (Smith et al. 2021), whereas the prelimbic cortex (PL) is engaged during cooperation for mutual rewards (Conde-Moro et al. 2024). On the other hand, the role of the infralimbic (IL) subregion in prosocial behavior has not been investigated. In addition, disrupting other cortical areas, such as the insular cortex (IC), decreased prosocial actions, including helping (Cox et al. 2022) and interacting with vulnerable conspecifics (Rogers-Carter et al. 2018), highlighting that prosocial decisions depend on coordinated activity across multiple cortical regions.

Consistent with this view, emerging evidence indicates that social decision-making also relies on interactions between cortical and subcortical circuits. For example, communication between PL and the basolateral amygdala (BLA) is required to share a reward with a partner (Scheggia et al. 2022), ACC-mediodorsal thalamus (MD) connectivity supports harm avoidance toward conspecifics (Song et al. 2023), and the PL-nucleus accumbens (NAc) oscillatory coherence increased during cooperation (Conde-Moro et al. 2024). Despite these advances, the broader cortical-subcortical networks that regulate the balance between prosocial and selfish behavior remain largely unexplored. In addition, most studies of social decision-making in mice have focused on C57BL/6 mice, an inbred strain with a well-characterized prosocial phenotype (Wu et al. 2021; Scheggia et al. 2022; Song et al. 2023; Pozo et al. 2023; Misiołek et al. 2023; Zhang et al. 2024), leaving open the question of whether these neural mechanisms generalize across genetically diverse populations.

To address these gaps, we examined neural substrates of social decision-making using inbred male C57BL/6 mice and outbred male CD1 mice. In addition to considering the regions mentioned above, namely mPFC, NAcC, MD, and BLA, we also considered the lateral hypothalamus (LH), implicated in social competition (Padilla-Coreano et al. 2022); the ventral tegmental area (VTA), which encodes the motivational value of social interactions (Hung et al. 2017); and the ventral hippocampus (vHPC), which processes social memory critical for guiding social decisions (Phillips et al. 2019). Here, using a social decision-making task combined with c-Fos mapping, we examined activity in a broad network during social decision-making in both inbred and outbred mouse strains. Our findings show that increased activity in the mPFC and nucleus accumbens core (NAcC) is associated with biases toward selfish choices in CD1 male mice. At the network level, social decision-making recruited coordinated activity between VTA, NAc, and the mPFC. However, network dynamics in prosocial vs selfish mice differed, with distinct prefrontal–subcortical network configurations recruited with BLA as a common hub.

## RESULTS

### Prosocial and selfish behaviors differ between strains of male mice

This study evaluated neural activity associated with prosocial and selfish behavior at a network level in two different strains of mice (C57BL/6 and CD1). To study prosocial and selfish behavior, we modified a recently published task for social decision-making wherein mice decide between a prosocial option that delivers a reward to a familiar partner and a selfish option that provides a reward only to themselves (Scheggia et al. 2022). To facilitate associative learning between decisions and social outcomes, we incorporated a conditioned stimulus (CS) prior to operant conditioning that signals what happens to the partner, and a forced decision block to ensure that subject mice sampled both options prior to freely choosing (**Figure 1**).

**Figure 1.**
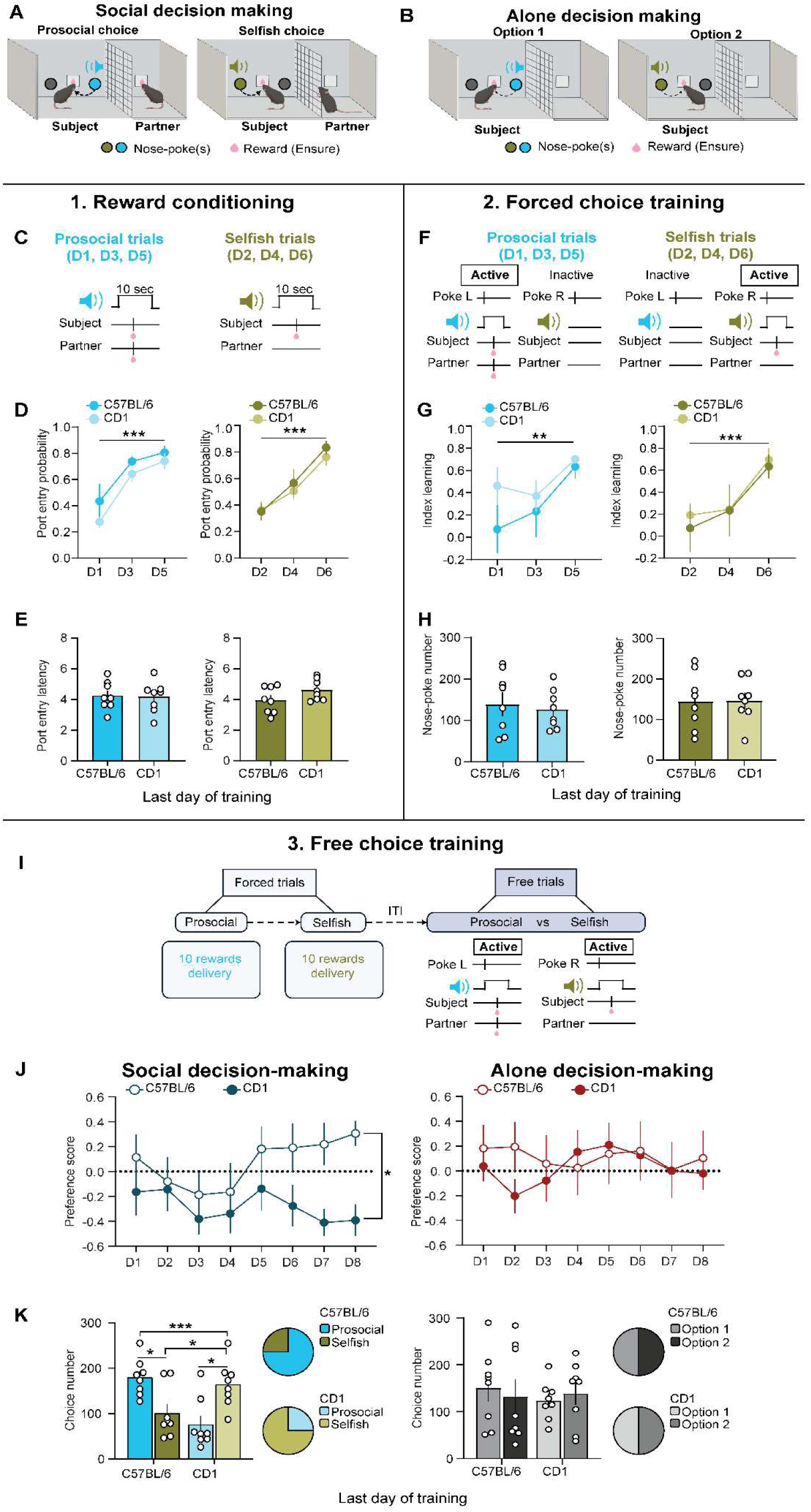
Strain differences in social decision-making. **A.** Social decision-making task. C57/BL6 and CD1 mice (n=8 per strain) choose between two nose-poke options: one that delivers a reward to themselves (selfish choice) and another that rewards them and a familiar partner (prosocial choice), who is a passive recipient (n=8 per strain). **B**. C57BL/6 (n=8) and CD1 (n=8) mice trained without a partner served as controls. **C.** Reward conditioning (stage1). Subject and partner mice received two distinct tones: one paired with rewards for both mice (prosocial trials) and another paired with rewards only for the subject mice (selfish trials). Prosocial (days 1, 3, and 5) and selfish (days 2, 4, and 6) trials were alternated every day. **D.** Reward port entries probability during prosocial and selfish trials in both C57BL/6 and CD1 mice across training days. **E.** Reward port entry latencies during prosocial and selfish trials in both C57BL/6 and CD1 mice at the end of training show learning over days. **F.** Forced choice training. Nose-poke responses were associated with prosocial or selfish outcomes. An active nose-poke triggered the previously conditioned tones, followed by reward delivery to both mice (prosocial trials) or to the subject only (selfish trials), whereas the inactive nose-poke had no consequences. Active and inactive nose-pokes were alternated daily across sessions (prosocial: days 1, 3, and 5; selfish: days 2, 4, and 6). **G.** Index learning shows discrimination between active and inactive nose-pokes during prosocial and selfish trials across days in both mouse strains. **H.** Nose-poke responses did not differ between mouse strains during prosocial or selfish trials at the end of training. **I.** Free choice training. After a block of forced trials (20 reward deliveries as a criterion), mice received free-choice trials with both nose-pokes activated. **J.** Left, preference score (prosocial – selfish choices/ sum of both) showed higher prosocial preference in C57BL/6 mice compared to CD1 mice during social decision-making. Right, preference score showed no side preference during alone decision-making. **K.** Left, choice number during the last day of training. C57BL/6 mice made more prosocial choices than selfish ones, whereas CD1 mice made more selfish than prosocial choices during social decision-making. Right, C57BL/6 and CD1 mice showed a similar number of Option 1 and Option 2 choices during alone decision-making. The pie chart showed the proportions of C57BL/6 and CD1 mice that made majority prosocial or selfish choices during social decision-making and during alone decision-making with Option 1 or Option 2. Data are mean ± SEM; *P < 0.05, **P < 0.01, ***P < 0.001.

Social decision-making task training (20 days) consisted of three consecutive stages: (1) reward conditioning training (**Figure 1C**), (2) forced-choice training (**Figure 1F**), and (3) free-choice training (**Figure 1I**). This protocol was designed to allow mice to express a social preference between two social outcomes after forced exploration of both options, and thus facilitate associations between actions and social outcomes, while minimizing side preferences formed by habit. To determine whether the presence of a partner guides choice behavior, an additional group of mice was trained under the same conditions but without a partner, serving as a control group (alone decision-making). In this condition, subjects decided between two options that delivered the same amount of reward but used the same two CS as our social group (**Figure 1B**).

During stage 1, reward conditioning, mice associated distinct tones with the prosocial or selfish outcomes (CS-prosocial or CS-selfish). Each CS was presented with a variable inter-trial-interval 30 times per day and alternated across days (CS-prosocial days 1, 3, and 5; CS-selfish 2, 4, and 6; **Figure 1C**). All subject mice showed successful conditioning to both CS as demonstrated by probability for port entries increasing over days (**Figure 1D**; two-way RM ANOVA; CS-prosocial: strain, F(1, 14) = 2.85, p = 0.11; days, F(2, 28) = 19.41, p < 0.001; interaction, F(2, 28) = 0.21, p = 0.80; CS-selfish: strain, F(1, 14) = 0.25, p = 0.62; days, F(2, 28) = 30.22, p < 0.001; interaction, F(2, 28) = 0.29, p = 0.74). Importantly, by the end of stage 1, reward port entry latency did not differ by strain (**Figure 1E**, unpaired-test: CS-prosocial, t(14) = 0.14, p= 0.88; CS-selfish, t(14) = 1.65, p= 0.11). Partner mice, those passively receiving rewards, also showed successful conditioning and discriminated between the CS-prosocial and CS-selfish, evidenced by increased port entries selectively to CS-prosocial (**Figure S1**; CS-prosocial: two-way RM ANOVA; strain, F(1,14)= 1.48, p=0.24; days, F(1.18, 16.73)=14.65, p= 0.0002; interaction, F (2, 28)= 0.29, p= 0.74; CS-selfish: two way RM ANOVA; strain, F(1,14)=1.81, p= 0.19, days, F(1.83, 25.64)= 0.13, p= 0.85; interaction, F(2, 28)=0.92, p= 0.40). This behavioral change in partners provides important additional social cues for subjects to learn the social outcomes of their choices. Finally, mice in the alone decision-making group (**Figure 1B**) showed successful acquisition of the CS-reward associations (**Figure S1B-C**).

In stage 2, forced choice training, mice were trained to associate nose-pokes with the prosocial or selfish CS and reward outcomes (**Figure 1F**). Every day, mice were presented with one nose poke that was active at a time, and nose-pokes elicited presentation of the CS-prosocial or the CS-selfish, followed by the corresponding reward outcomes. Using this configuration, mice were forced to use one nose-poke per session. All mice showed successful discrimination of active versus inactive nose-pokes across days (two-way RM ANOVA; prosocial: strain, F(1, 14)= 1.09, p=0.31; days, F(1.89,26,64)= 7.26, p=0.003; interaction, F(2, 28)= 1.052, p= 0.36; selfish: strain, F(1, 14)= 0.08, p=0.7; days, F(1.94,27,19)= 14.92, p<0.001; interaction, F(2, 28)= 0.14, p=0.86). Neither strain showed nose-poke preferences in this forced stage (**Figure 1H**; unpaired t-test: prosocial, t(14) = 0.66, p= 0.51; selfish, t(14) = 0.10, p= 0.91). Mice in the alone decision-making group also discriminated between active and inactive nose-pokes across days **(Figure S1D**; **Supp Figure 1E)**. These results indicate comparable discrimination between active and inactive nose-pokes across strains, demonstrating that both C57BL/6 and CD1 mice learned the operant associations.

Finally, in stage 3, free-choice training, we measured free-choice preferences between prosocial and selfish outcomes. Each session started with counterbalanced forced trials to ensure mice sampled both options, followed by 30 minutes with both nose-pokes active to evaluate choice preference (**Figure 1I**). We found that, across days, C57BL/6 mice developed a prosocial choice preference while CD1 mice developed a selfish preference (**Figure 1J**; two-way RM ANOVA; strain, F(1, 14)= 4.66, p=0.04; days, F(4.08, 57.8)= 1.3, p=0.26; interaction, (7, 98)= 1.42, p= 0.20). In contrast, alone decision-making resulted in no side preferences (**Figure 1J**; two-way RM ANOVA; strain, F(1, 14)= 0.12, p=0.73; days, F(4.09, 57.32)= 0.81; interaction, F(7, 98)= 1.057, p= 0.39). In the last day of training, C57BL/6 mice showed significantly more prosocial than selfish choices, whereas CD1 mice exhibited the opposite pattern (**Figure 1K**; two-way RM ANOVA; strain, F (1,14)=2.21, p=0.15; nose-poke, F(1,14)= 0.39, p=0.95; interaction (1,14)= 15.08, p= 0.001; Post-hoc comparisons: prosocial choice C57BL/6 vs CD1, p= 0.01; selfish choice C57BL/6 vs CD1, p= 0.0007; C57BL/6 prosocial vs selfish, p= 0.02; CD1 prosocial vs selfish, p= 0.02). In contrast, during alone decision-making, there was no choice preference (**Figure 1K**; two-way RM ANOVA; strain, F(1, 14)= 0.41; nose-poke, F(1,14)= 0.001, p=0.96; interaction, F (1,14)= 0.23, p= 0.63). Choice percentages did not differ across nose-poke sides or auditory cues on the final training day in either strain (two-way RM ANOVA; all p > 0.5), indicating that strain differences in social preference were driven by partner presence rather than task-related variables (**Figure S2A–B**). Together, our results confirm previous evidence of prosocial behavior in C57BL/6 mice (Scheggia et al. 2022; Song et al. 2023) while uncovering new findings about strain differences and antisocial preferences in CD1 male mice.

### mPFC activity correlates with social choices

Given the strain differences in social decision-making, we next examined whether brain activity patterns explained differences in social decision-making. Once mice showed a stable preference for selfish or prosocial choices, C57BL/6 and CD1 mice were tested on an additional day of social or alone decision-making (**Figure 2A-B**). To evaluate brain activity patterns, we labeled c-Fos expression immunohistochemically and compared the mean number of c-Fos+ cells in cortical regions implicated in prosocial behaviors, specifically prefrontal and insular cortices (Conde-Moro et al. 2024; Yamagishi et al. 2020; Song et al. 2023; Cox et al. 2022; Rogers-Carter et al. 2018).

**Figure 2.**
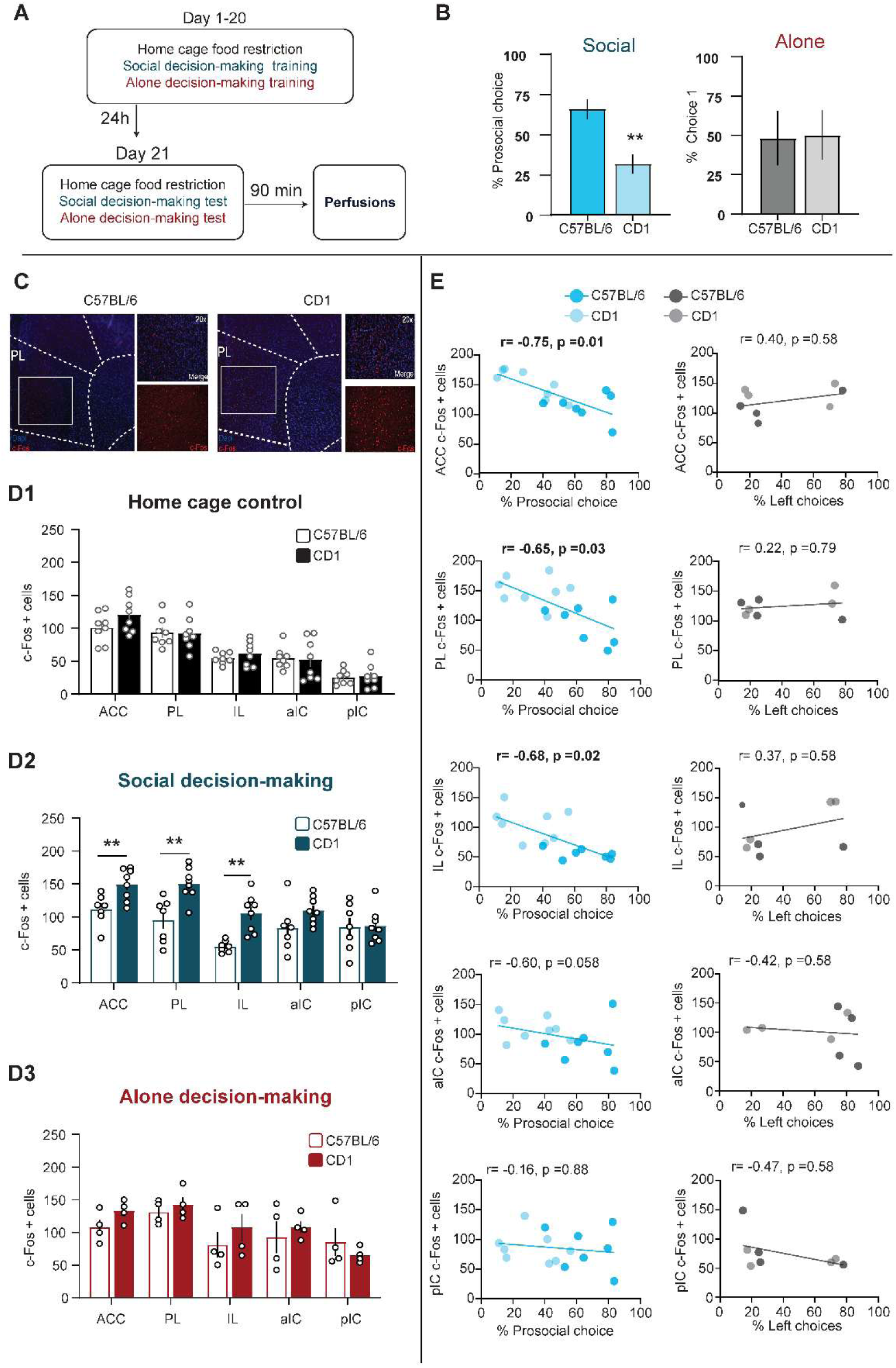
Prefrontal c-Fos activity is associated with selfish behavior. **A.** Experimental timeline. C57BL/6 and CD1 mice were tested in one of three conditions: home-cage food restriction (baseline c-Fos), social decision-making, or alone decision-making. After 21 days, mice were perfused for c-Fos mapping. **B.** Percentage of prosocial choice during social and alone decision making in mice prior to c-Fos mapping. **C.** Representative c-Fos images for PL of C57BL/6 and CD1 mice during social decision-making. **D.** Mean c-Fos positive cells count across ACC, PL, IL, aIC, and pIC of C57 and CD1 mice under home cage (**D1**; n=8 per strain), social (**D2**; c57BL/6 n=7; CD1, n=8), and alone decision-making (**D3**; n= 4 per strain). **E**. Top, relationship between cortical subregion c-Fos and percentage of prosocial choices during social decision-making or percentage of left choice during alone decision-making. P values were adjusted using a Benjamini–Hochberg FDR procedure.

Importantly, in a naïve baseline home cage condition there were no strain differences in c-Fos expression in mPFC subregions (ACC, PL, and IL), anterior IC (aIC), and posterior IC (pIC) (**Figure 2D1**; two-way RM ANOVA; strain, F(1, 14)= 0.63, p=0.44; regions, F(2.44, 36.24)= 48.72, p= <0.0001; interaction, F (4,56)= 0.84, p= 0.50). On the other hand, c-Fos expression during social decision-making differed across strains. CD1 mice had higher number of c-Fos+ cells in the ACC, PL, and IL compared with C57BL/6 mice (**Figure 2D2**; two-way RM ANOVA; strain, F (1,13)= 16.70, p= 0.001; regions, F(2.89,3758)= 13.06, p< 0.0001; interaction (4,52)= 2.29, p= 0.02; post hoc comparison, p= 0.01). Importantly, these differences were not observed during alone decision-making, indicating that specifically social decision-making recruited more prefrontal activity in CD1 compared to C57BL/6 mice (**Figure 2D3**; two-way RM ANOVA; strain, F(1, 6)= 0.68, p=0.44; regions, F(3.02, 18.12)= 6.32, p= 0.004; interaction, F (4,24)=0.98, p= 0.43).

Given that increased prefrontal activity was not due to overall mouse strain differences, we hypothesized that these differences were due to their social choices. To test this hypothesis, we correlated the number of c-Fos+ cells with the percentage of prosocial choices (**Figure 2E**). c-Fos activity was anticorrelated with the percentage of prosocial choices in ACC (r= -0.75, p= 0.014), PL (r= -0.65, p= 0.03), and IL (r=-0.68, p= 0.02). Because animals tested in the alone condition had no real prosocial option, we instead examined whether the percentage of left nose-poke choices correlated with c-Fos expression to rule out a side preference. No significant correlations were observed in the alone condition, indicating that the correlations observed in the social decision-making group are specific to the social context (**Figure 2E**).

### Strain-specific increases in NAcC c-Fos activity during social decision-making

We next examined c-Fos levels in select subcortical regions associated with social behaviors (Conde-Moro et al. 2024; Padilla-Coreano et al. 2022; Hung et al. 2017; Phillips et al. 2019; Scheggia et al. 2022) in a naïve baseline home cage group, social, and alone decision-making groups of both strains of mice. After baseline home-cage food-restriction conditions, there was no significant effect of strain in c-Fos+ cells in subcortical regions (**Figure 3C1**; two-way RM ANOVA; strain, F (1,14)= 2.15, p= 0.16; regions, F(3.96, 55.45)= 22.48, p< 0.0001; interaction (6,84)= 0.82, p= 0.55). However, c-Fos activity during social decision-making showed a significant interaction between strain and region, with higher NAcC c-Fos expression in CD1 mice (**Figure 3C2**; two-way RM ANOVA; strain, F (1,13)= 0.44, p= 0.51; regions, F(3.70,41.21)= 11.12, p<0.0001; interaction (6,78)= 7.94, p< 0.0001; post hoc comparison, p= 0.001). On the other hand, c-Fos activity during alone decision-making also showed a significant interaction between strain and region, yet no specific region had significant differences between strains (**Figure 3C3;** two-way RM ANOVA; strain, F (1,6)= 0.002, p= 0.95; regions, F(2.84,17.09)= 11.83, p< 0.0002; interaction (6,36)= 2.42, p= 0.04; NAcC, post hoc comparison, p= 0.09). These results suggest that NAcC activity could be driven by more selfish choices in CD1 mice. NAcC cFos+ cells and the percentage of prosocial choices showed a trending anti-correlation (**Figure 3D**), suggesting that social choices may influence c-Fos expression in NAcC during social decision making in CD1 mice.

**Figure 3.**
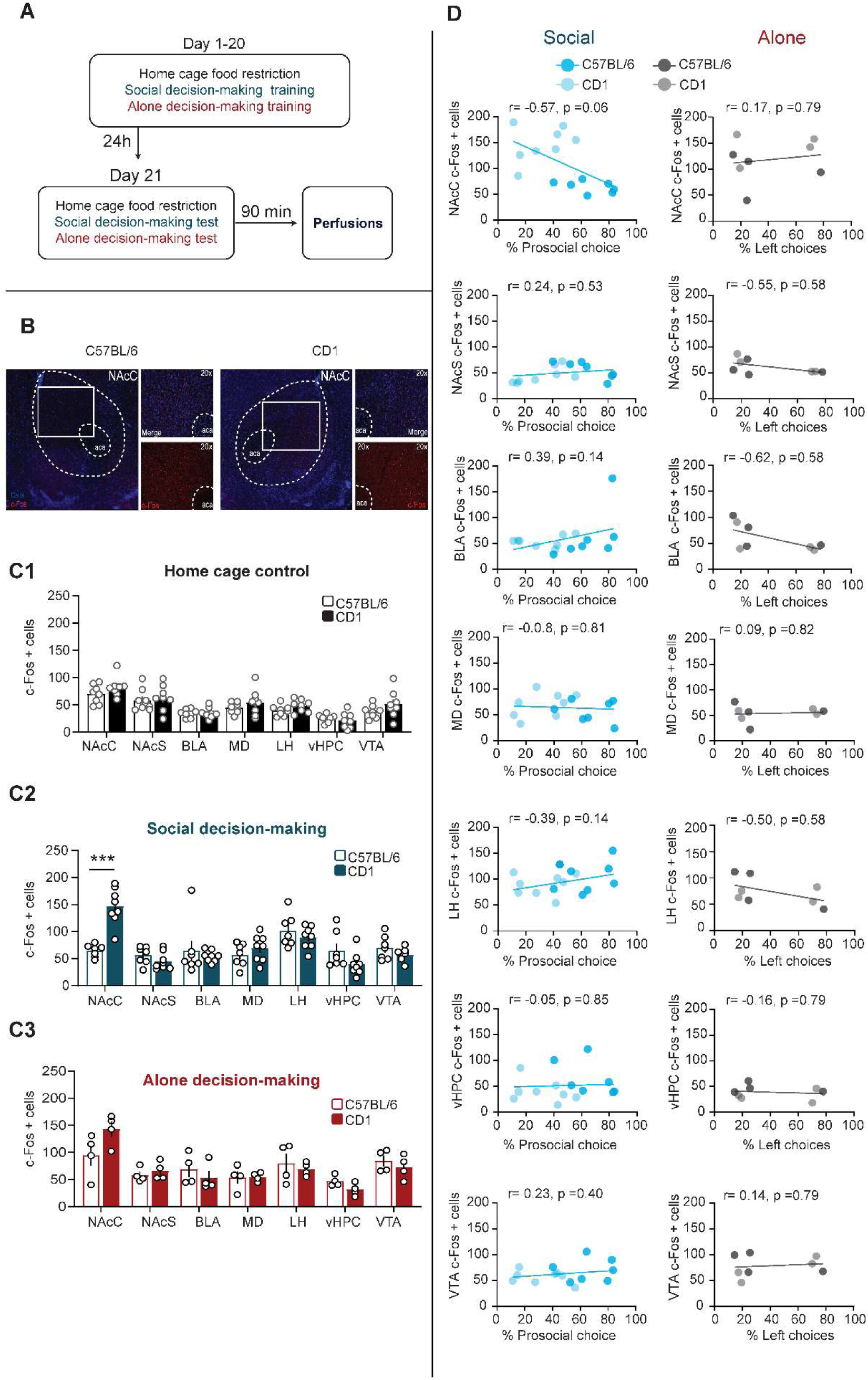
Elevated NAcC c-Fos activation in selfish CD1 mice. **A.** Experimental timeline. **B**. Representative c-Fos images for C57BL/6 and CD1 mice for NacC during social decision-making. **C.** Mean c-Fos positive cells count across NAcC, NAcS, BLA, MD, LH, vHPC, and VTA of C57 and CD1 mice under (**C1)** home cage, (**C2**) social, and (**C3**) alone decision-making. **D**. Top, Relationship between subcortical c-Fos expression and the percentage of prosocial choices during social decision-making or left choice during alone decision-making. P values were adjusted using a Benjamini–Hochberg FDR procedure.

Given that mPFC activity and NAcC are associated with reward processing (Burgos-Robles et al. 2013; Soares-Cunha et al. 2020) and social interactions (Lee et al. 2016; Dai et al. 2022), we evaluated whether c-Fos activity during social decision-making was influenced by reward and social interactions. No significant correlations were observed between c-Fos expression and the total number of rewards consumed during the social decision-making (**Figure S3A**), nor the alone decision-making (**Figure S3B**). Despite significant differences in the number of face-to-face social interactions across strains of mice (**Figure S4A**), the total number of these bouts did not correlate with cortical or subcortical c-Fos expression (**Figure S4B**), indicating that c-Fos activity was not related to general reward consumption or social engagement. Overall, these results reinforce that these activity patterns are driven by differences in social choices. Finally, c-Fos expression was not correlated with side choices (percentage of left nose-poke responses) or tone-guided choices (percentage of Tone 1 responses) in the social condition (**Figure S5**), and no significant correlations were observed with side or tone-guided choices in the alone condition (**Figure S6**).

### Network states and cross-regional correlations depend on social decisions

To investigate how interactions across regions relate to social decision-making, we next examined functional connectivity among these cortical and subcortical regions. Because baseline c-Fos expression did not differ between C57BL/6 and CD1 mice (**Figure 2D and 3C**) and to assess networks recruited for general social decision-making, data from both strains was combined for cross-correlation analyses of c-Fos+ cells across brain regions. Functional connectivity was assessed first by using Network-Based Statistics (NBS; 5,000 permutations; primary threshold p <0.01; Zalesky et al. 2010), as social decision-making is thought to engage distributed circuits rather than isolated regional interactions (Rogers-Carter and Christianson 2019). Because NBS is used to identify network interactions (i.e., three or more interconnected brain regions), we additionally examined individual pairwise relationships. We used a permutation test to determine which pairwise cross-correlations of c-Fos+ cell counts across brain region pairs exceeded chance level as previously described (De Paula Cunha Almeida et al. 2025).

During social decision-making, a significant connected subnetwork component (extent = 3 edges, p = 0.0068) comprising the VTA, NAcC, PL, and IL was found (**Figure 4A**, **Table 1**). These results indicate coordinated subnetwork activity within a corticostriatal–midbrain subnetwork during social decision-making, supporting functional coupling among prefrontal (PL, IL), striatal (NAcC), and midbrain (VTA) regions. This was further supported by pairwise positive correlations that exceeded chance levels between mPFC subregions, NAcC, and VTA (PL-IL, r = 0.6790, p=0.0029; NAcC–PL, r = 0.7065, p=0.0019; NAcC-IL, r=0.6022, p=0.0229; VTA-NAcC, r= 0.6523, p=0.0089). Importantly, none of these cross-correlations were significant in the alone decision-making group, and NBS found no significant subnetworks in this group (**Figure S7; Table 2**). We also observed more pairwise correlations that exceeded chance levels in social decision-making (13 pairwise relationships; **Figure 4A**) than in the alone condition (3 pairs; **Figure S7A; Table 2**). Altogether, these findings suggest that coordinated corticostriatal–midbrain coupling is specifically recruited to enable social decision-making.

**Figure 4.**
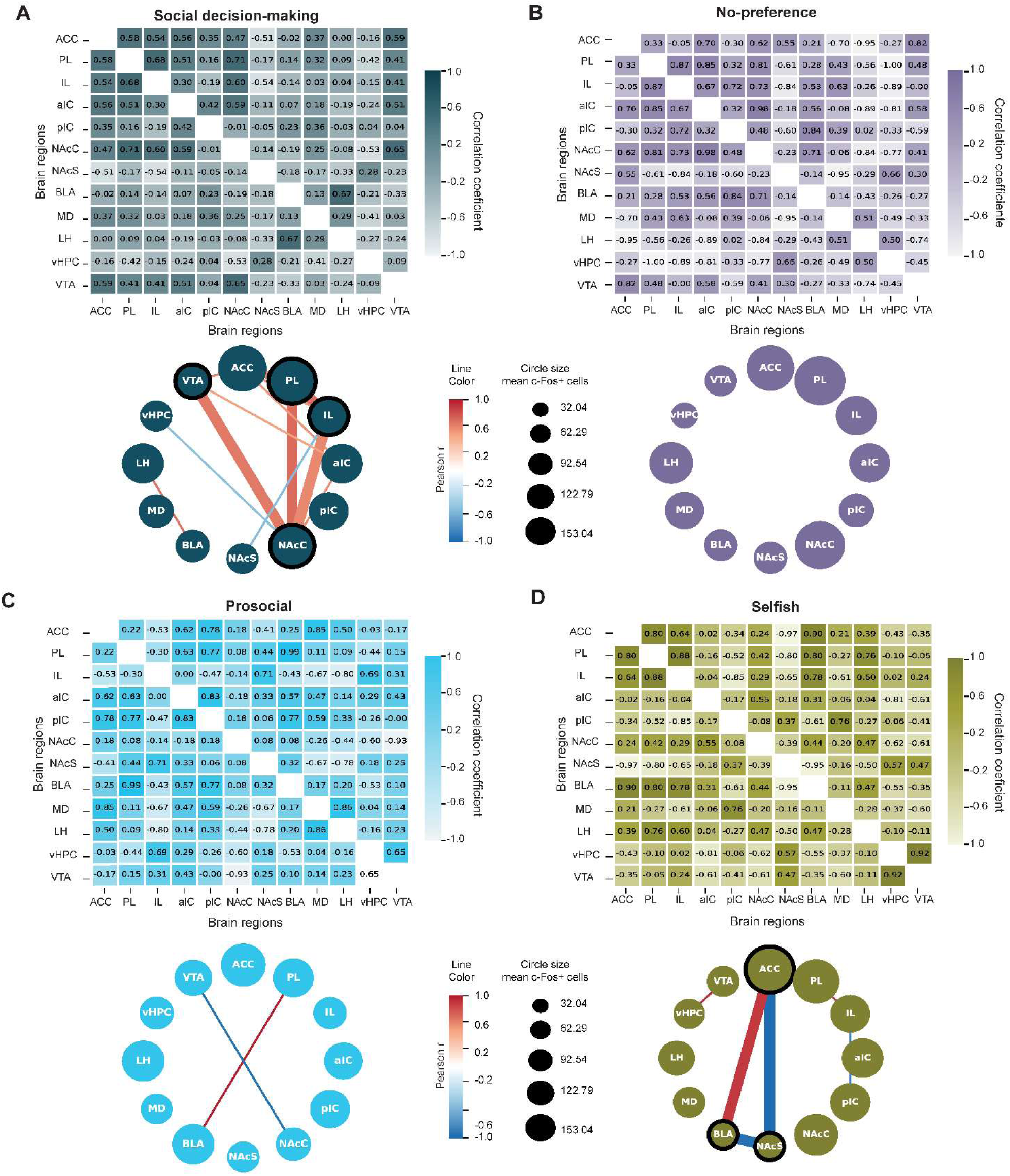
Cross-regional c-Fos correlations across social decision-making conditions. **A–D**, Heatmaps (top) showing pairwise Pearson correlations of c-Fos+ cell counts across brain regions and corresponding network representations (bottom) illustrating suprathreshold cross-regional associations during social decision-making for all mice (**A**), prosocial mice (**B**), selfish mice (**C**), and mice with no preference (**D**). Network visualizations show significant cross-correlations for each condition based on a null-distribution method. Line color represents Pearson r value. Only cross-correlations in the top/bottom 2.5% of null distribution and p<0.05 in permutation test are shown. Black circle outlines and thick lines indicate significant components through NBS (p<0.01). The circle size represents the mean c-Fos expression for each region. Sample sizes for social decision-making (n=8 C57 and 8 CD1), prosocial mice (n=5 C57), selfish (n=5 CD1 and 1 C57), and no-preference (n= 3 CD1 and 1 C57).

**Table 1.** Pearson’s correlations of mean c-Fos + cells count across brain regions during social decision-making.

| Pair region | Pearson's R | Permutation<br>*p*-value | Adjusted<br>*p*-value (FDR) | Percentile of<br>observed<br>value in the<br>null<br>distribution | n |
| --- | --- | --- | --- | --- | --- |
| **ACC-PL** | **0.581949898** | **0.01999001** | **0.208895552** | **99.05** | **15** |
| **ACC-IL** | **0.543343631** | **0.03798101** | **0.208895552** | **98.15** | **15** |
| **ACC-aIC** | **0.556415301** | **0.03598201** | **0.208895552** | **98.25** | **15** |
| ACC-pIC | 0.352968271 | 0.19990005 | 0.527736132 | 90.05 | 15 |
| ACC-NAcC | 0.474069113 | 0.05997002 | 0.247376312 | 97.05 | 15 |
| **ACC-NAcS** | **-0.51446661** | **0.04897551** | **0.228685657** | **2.4** | **15** |
| ACC-BLA | -0.02334483 | 0.97651174 | 0.976511744 | 48.8 | 15 |
| ACC-MD | 0.373511449 | 0.17491254 | 0.501922952 | 91.3 | 15 |
| ACC-LH | 0.002540277 | 0.94452774 | 0.974044228 | 52.8 | 15 |
| ACC-vHPC | -0.158109 | 0.54772614 | 0.83958021 | 27.35 | 15 |
| **ACC-VTA** | **0.586379709** | **0.0129935** | **0.208895552** | **99.4** | **15** |
| **PL-IL** | **0.67885679** | **0.0029985** | **0.098950525** | **99.9** | **15** |
| PL-aIC | 0.50856697 | 0.05097451 | 0.228685657 | 97.5 | 15 |
| PL-pIC | 0.156826789 | 0.55772114 | 0.83958021 | 72.15 | 15 |
| **PL-NAcC** | **0.706528415** | **0.001999** | **0.098950525** | **99.95** | **15** |
| PL-NAcS | -0.1669304 | 0.53573213 | 0.83958021 | 26.75 | 15 |
| PL-BLA | 0.135544302 | 0.64567716 | 0.887806097 | 67.75 | 15 |
| PL-MD | 0.32212734 | 0.22388806 | 0.568331219 | 88.85 | 15 |
| PL-LH | 0.086148421 | 0.7876062 | 0.928250161 | 60.65 | 15 |
| PL-vHPC | -0.42340361 | 0.12793603 | 0.425487256 | 6.35 | 15 |
| PL-VTA | 0.406713747 | 0.12893553 | 0.425487256 | 93.6 | 15 |
| IL-aIC | 0.297373859 | 0.28485757 | 0.66605407 | 85.8 | 15 |
| IL-pIC | -0.19299457 | 0.50874563 | 0.839430285 | 25.4 | 15 |
| **IL-NAcC** | **0.602268296** | **0.02298851** | **0.208895552** | **98.9** | **15** |
| **IL-NAcS** | **-0.54227003** | **0.03298351** | **0.208895552** | **1.6** | **15** |
| IL-BLA | -0.13995899 | 0.7836082 | 0.928250161 | 39.15 | 15 |
| IL-MD | 0.028243553 | 0.91554223 | 0.960186573 | 54.25 | 15 |
| IL-LH | 0.04337691 | 0.83858071 | 0.954247014 | 58.1 | 15 |
| IL-vHPC | -0.15067607 | 0.62568716 | 0.887806097 | 31.25 | 15 |
| IL-VTA | 0.406096984 | 0.14992504 | 0.471192975 | 92.55 | 15 |
| aIC-pIC | 0.415085022 | 0.12093953 | 0.425487256 | 94 | 15 |
| **aIC-NAcC** | **0.588739836** | **0.01999001** | **0.208895552** | **99.05** | **15** |
| aIC-NAcS | -0.11291904 | 0.71564218 | 0.926125173 | 35.75 | 15 |
| aIC-BLA | 0.069387369 | 0.75362319 | 0.928250161 | 62.35 | 15 |
| aIC-MD | 0.178885951 | 0.50674663 | 0.839430285 | 74.7 | 15 |
| aIC-LH | -0.19275765 | 0.44377811 | 0.770772508 | 22.15 | 15 |
| aIC-vHPC | -0.24298591 | 0.3948026 | 0.743466105 | 19.7 | 15 |
| aIC-VTA | 0.511406291 | 0.05197401 | 0.228685657 | 97.45 | 15 |
| pIC-NAcC | -0.01489024 | 0.97251374 | 0.976511744 | 48.6 | 15 |
| pIC-NAcS | -0.0505559 | 0.83458271 | 0.954247014 | 41.7 | 15 |
| pIC-BLA | 0.230284807 | 0.4067966 | 0.743466105 | 79.7 | 15 |
| pIC-MD | 0.356219862 | 0.18990505 | 0.522238881 | 90.55 | 15 |
| pIC-LH | -0.03332337 | 0.91654173 | 0.960186573 | 45.8 | 15 |
| pIC-vHPC | 0.03615425 | 0.87056472 | 0.960186573 | 56.5 | 15 |
| pIC-VTA | 0.044077866 | 0.90654673 | 0.960186573 | 54.7 | 15 |
| NAcC-NAcS | -0.13961707 | 0.58770615 | 0.861969015 | 29.35 | 15 |
| NAcC-BLA | -0.18815136 | 0.69265367 | 0.926125173 | 34.6 | 15 |
| NAcC-MD | 0.249804227 | 0.35682159 | 0.727636182 | 82.2 | 15 |
| NAcC-LH | -0.08298228 | 0.7856072 | 0.928250161 | 39.25 | 15 |
| **NAcC-vHPC** | **-0.52682354** | **0.03698151** | **0.208895552** | **1.8** | **15** |
| **NAcC-VTA** | **0.652379075** | **0.0089955** | **0.197901049** | **99.6** | **15** |
| NAcS-BLA | -0.1777702 | 0.64367816 | 0.887806097 | 32.15 | 15 |
| NAcS-MD | -0.16574212 | 0.55972014 | 0.83958021 | 27.95 | 15 |
| NAcS-LH | -0.32949241 | 0.24387806 | 0.596146371 | 12.15 | 15 |
| NAcS-vHPC | 0.281692252 | 0.31284358 | 0.66605407 | 84.4 | 15 |
| NAcS-VTA | -0.22922223 | 0.4167916 | 0.743466105 | 20.8 | 15 |
| BLA-MD | 0.12571286 | 0.71464268 | 0.926125173 | 64.3 | 15 |
| **BLA-LH** | **0.674444304** | **0.03198401** | **0.208895552** | **98.45** | **15** |
| BLA-vHPC | -0.21103092 | 0.4007996 | 0.743466105 | 20 | 15 |
| BLA-VTA | -0.33123579 | 0.16191904 | 0.485757121 | 8.05 | 15 |
| MD-LH | 0.291701801 | 0.30684658 | 0.66605407 | 84.7 | 15 |
| MD-vHPC | -0.40706297 | 0.12593703 | 0.425487256 | 6.25 | 15 |
| MD-VTA | 0.030620477 | 0.90054973 | 0.960186573 | 55 | 15 |
| LH-vHPC | -0.27440574 | 0.31084458 | 0.66605407 | 15.5 | 15 |
| LH-VTA | -0.24351178 | 0.36381809 | 0.727636182 | 18.15 | 15 |
| vHPC-VTA | -0.08886915 | 0.7846077 | 0.928250161 | 39.2 | 15 |

Adjusted p-values were obtained from permutation-derived null distributions of Pearson correlation coefficients and corrected for multiple comparisons using the Benjamini–Hochberg false discovery rate (FDR). Percentile values indicate the position of each observed correlation relative to the permutation-based null distribution. Bold rows indicate correlations falling within the upper or lower 2.5% of the null distribution and with permutation $p < 0.05$ (uncorrected). $n$ indicates the number of animals included in each correlation analysis.

**Table 2.** Pearson’s correlations of mean c-Fos + cells count across brain regions during alone decision-making.

| Pair region | Pearson's R | Permutation *p*-value | Adjusted *p*-value (FDR) | Percentile of observed value in the null distribution | n |
| --- | --- | --- | --- | --- | --- |
| ACC-PL | 0.045413914 | 0.943528236 | 0.988458 | 52.85 | 8 |
| ACC-IL | 0.242573136 | 0.60169915 | 0.891154 | 69.95 | 8 |
| ACC-aIC | -0.01156472 | 0.990504748 | 0.990505 | 49.5 | 8 |
| ACC-pIC | -0.1318678 | 0.785607196 | 0.925894 | 39.25 | 8 |
| ACC-NAcC | 0.450295652 | 0.286856572 | 0.891154 | 85.7 | 8 |
| ACC-NAcS | 0.224724944 | 0.604697651 | 0.891154 | 69.8 | 8 |
| ACC-BLA | -0.27964017 | 0.498750625 | 0.891154 | 24.9 | 8 |
| ACC-MD | 0.350436814 | 0.410794603 | 0.891154 | 79.5 | 8 |
| ACC-LH | -0.40156532 | 0.335832084 | 0.891154 | 16.75 | 8 |
| ACC-vHPC | -0.25568887 | 0.526736632 | 0.891154 | 26.3 | 8 |
| ACC-VTA | -0.33991422 | 0.396801599 | 0.891154 | 19.8 | 8 |
| PL-IL | 0.631301432 | 0.124937531 | 0.775112 | 93.8 | 8 |
| PL-aIC | 0.505515368 | 0.187906047 | 0.775112 | 90.65 | 8 |
| PL-pIC | 0.078234353 | 0.831584208 | 0.944782 | 58.45 | 8 |
| PL-NAcC | 0.527294147 | 0.177911044 | 0.775112 | 91.15 | 8 |
| PL-NAcS | -0.51636538 | 0.15992004 | 0.775112 | 7.95 | 8 |
| PL-BLA | -0.04758013 | 0.971514243 | 0.990505 | 48.55 | 8 |
| PL-MD | -0.1886315 | 0.585707146 | 0.891154 | 29.25 | 8 |
| PL-LH | 0.59586216 | 0.151924038 | 0.775112 | 92.45 | 8 |
| PL-vHPC | -0.00643244 | 0.978510745 | 0.990505 | 48.9 | 8 |
| PL-VTA | 0.706407119 | 0.045977011 | 0.775112 | 97.75 | 8 |
| IL-aIC | 0.252035803 | 0.544727636 | 0.891154 | 72.8 | 8 |
| IL-pIC | 0.378668657 | 0.52173913 | 0.891154 | 73.95 | 8 |
| IL-NAcC | 0.446561097 | 0.261869065 | 0.891154 | 86.95 | 8 |
| IL-NAcS | -0.35822793 | 0.374812594 | 0.891154 | 18.7 | 8 |
| IL-BLA | -0.11464297 | 0.724637681 | 0.891154 | 36.2 | 8 |
| IL-MD | 0.576001068 | 0.111944028 | 0.775112 | 94.45 | 8 |
| IL-LH | 0.171764934 | 0.667666167 | 0.891154 | 66.65 | 8 |
| IL-vHPC | -0.33396607 | 0.446776612 | 0.891154 | 22.3 | 8 |
| IL-VTA | 0.416277593 | 0.289855072 | 0.891154 | 85.55 | 8 |
| aIC-pIC | 0.382505567 | 0.357821089 | 0.891154 | 82.15 | 8 |
| aIC-NAcC | 0.545496544 | 0.182908546 | 0.775112 | 90.9 | 8 |
| aIC-NAcS | -0.18113222 | 0.707646177 | 0.891154 | 35.35 | 8 |
| aIC-BLA | 0.505056795 | 0.178910545 | 0.775112 | 91.1 | 8 |
| aIC-MD | -0.13190477 | 0.84057971 | 0.944782 | 42 | 8 |
| **aIC-LH** | **0.801177812** | **0.011994003** | **0.775112** | **99.45** | **8** |
| aIC-vHPC | -0.38207164 | 0.354822589 | 0.891154 | 17.65 | 8 |
| aIC-VTA | 0.32199103 | 0.468765617 | 0.891154 | 76.6 | 8 |
| pIC-NAcC | 0.110848154 | 0.844577711 | 0.944782 | 57.8 | 8 |
| pIC-NAcS | 0.056631978 | 0.741629185 | 0.891154 | 62.95 | 8 |
| **pIC-BLA** | **0.727138841** | **0.030984508** | **0.775112** | **98.5** | **8** |
| pIC-MD | 0.63362717 | 0.056971514 | 0.775112 | 97.2 | 8 |
| pIC-LH | 0.531188246 | 0.263868066 | 0.891154 | 86.85 | 8 |
| pIC-vHPC | 0.127789943 | 0.736631684 | 0.891154 | 63.2 | 8 |
| pIC-VTA | 0.387476549 | 0.429785107 | 0.891154 | 78.55 | 8 |
| NAcC-NAcS | -0.10763719 | 0.742628686 | 0.891154 | 37.1 | 8 |
| NAcC-BLA | 0.31615185 | 0.485757121 | 0.891154 | 75.75 | 8 |
| NAcC-MD | 0.12444853 | 0.700649675 | 0.891154 | 65 | 8 |
| NAcC-LH | 0.227709009 | 0.603698151 | 0.891154 | 69.85 | 8 |
| NAcC-vHPC | -0.56363248 | 0.150924538 | 0.775112 | 7.5 | 8 |
| NAcC-VTA | 0.360059097 | 0.394802599 | 0.891154 | 80.3 | 8 |
| NAcS-BLA | 0.107428751 | 0.734632684 | 0.891154 | 63.3 | 8 |
| NAcS-MD | 0.194671269 | 0.616691654 | 0.891154 | 69.2 | 8 |
| NAcS-LH | -0.32403511 | 0.435782109 | 0.891154 | 21.75 | 8 |
| NAcS-vHPC | 0.0512625 | 0.898550725 | 0.96397 | 55.1 | 8 |
| NAcS-VTA | -0.67659103 | 0.086956522 | 0.775112 | 4.3 | 8 |
| BLA-MD | 0.165057426 | 0.72163918 | 0.891154 | 63.95 | 8 |
| BLA-LH | 0.590067421 | 0.134932534 | 0.775112 | 93.3 | 8 |
| BLA-vHPC | 0.01857546 | 0.897551224 | 0.96397 | 55.15 | 8 |
| BLA-VTA | 0.404743882 | 0.295852074 | 0.891154 | 85.25 | 8 |
| MD-LH | -0.24515489 | 0.540729635 | 0.891154 | 27 | 8 |
| MD-vHPC | -0.20480099 | 0.614692654 | 0.891154 | 30.7 | 8 |
| MD-VTA | -0.06720861 | 0.905547226 | 0.96397 | 45.25 | 8 |
| LH-vHPC | 0.159517564 | 0.68165917 | 0.891154 | 65.95 | 8 |
| LH-VTA | 0.657736234 | 0.071964018 | 0.775112 | 96.45 | 8 |
| vHPC-VTA | 0.220192255 | 0.595702149 | 0.891154 | 70.25 | 8 |

Adjusted p-values were obtained from permutation-derived null distributions of Pearson correlation coefficients and corrected for multiple comparisons using the Benjamini–Hochberg false discovery rate (FDR). Percentile values indicate the position of each observed correlation relative to the permutation-based null distribution. Bold rows indicate correlations falling within the upper or lower 2.5% of the null distribution and with permutation $p < 0.05$ (uncorrected). $n$ indicates the number of animals included in each correlation analysis.

We next evaluated whether social choices influenced the network state. We separated our subject population according to their overall preferred social choice on the last day of training and grouped them into prosocial, selfish, and no-preference mice. Mice with no social preference had no significant pairwise correlation nor subnetwork activity (**Figure 4B**, **Table 3)**. On the other hand, mice with clear social preferences showed distinct network interactions. Prosocial mice had two pairwise correlations exceeding chance level with PL–BLA showing a strong positive correlation (**Figure 4C**, **Table 4**, r = 0.9895, p=0.0289) and NAcC-VTA showing a strong negative correlation (**Figure 4C**, **Table 4**, r = -0.9326, p=0.0359). In contrast, selfish mice exhibited a significant connected component (NBS, extent = 2 edges, p= 0.041) comprising ACC, NAcS, and BLA (**Figure 4D**, **Table 5**), indicating that network interactions are altered depending on social preferences. Consistent with this subnetwork, the permutation test identified a strong correlation between ACC-BLA and anticorrelations between NAcS-BLA and NAcS-ACC (NAcS–BLA r = −0.9496, p=0.009; ACC-BLA r = 0.8980, p=0.046; ACC–NAcS, r = −0.9720, p=0.049; **Figure 4D**, **Table 5**). Our results identify a novel subnetwork of coordinated activity in ACC–NAcS–BLA during antisocial behavior and confirm previous findings identifying the PL-BLA circuit as mediating prosocial behavior.

**Table 3.** Pearson’s correlations of mean c-Fos + cells count across brain regions during no-preference choices.

| **Pair region** | **Pearson's R** | **Permutation**<br>***p*-value** | **Adjusted**<br>***p*-value**<br>**(FDR)** | **Percentile of**<br>**observed**<br>**value in the**<br>**null**<br>**distribution** | **n** |
| --- | --- | --- | --- | --- | --- |
| ACC-PL | 0.325951 | 0.666667 | 0.862745 | 68.83681 | 4 |
| ACC-IL | -0.04693 | 0.833333 | 0.932203 | 41.05903 | 4 |
| ACC-aIC | 0.704478 | 0.25 | 0.862745 | 89.67014 | 4 |
| ACC-pIC | -0.30012 | 0.75 | 0.868421 | 35.50347 | 4 |
| ACC-NAcC | 0.618089 | 0.333333 | 0.862745 | 84.80903 | 4 |
| ACC-NAcS | 0.545785 | 0.416667 | 0.862745 | 81.77083 | 4 |
| ACC-BLA | 0.214917 | 0.75 | 0.868421 | 66.05903 | 4 |
| ACC-MD | -0.70034 | 0.25 | 0.862745 | 11.02431 | 4 |
| ACC-LH | -0.94845 | 0.166667 | 0.862745 | 6.336806 | 4 |
| ACC-vHPC | -0.26876 | 0.833333 | 0.932203 | 40.45139 | 4 |
| ACC-VTA | 0.818669 | 0.25 | 0.862745 | 89.67014 | 4 |
| PL-IL | 0.871141 | 0.166667 | 0.862745 | 93.83681 | 4 |
| PL-aIC | 0.845755 | 0.333333 | 0.862745 | 86.19792 | 4 |
| PL-pIC | 0.321987 | 0.75 | 0.868421 | 64.84375 | 4 |
| PL-NAcC | 0.805425 | 0.333333 | 0.862745 | 86.19792 | 4 |
| PL-NAcS | -0.61063 | 0.5 | 0.862745 | 22.30903 | 4 |
| PL-BLA | 0.280425 | 0.75 | 0.868421 | 66.05903 | 4 |
| PL-MD | 0.430932 | 0.666667 | 0.862745 | 69.01042 | 4 |
| PL-LH | -0.55582 | 0.5 | 0.862745 | 23.00347 | 4 |
| PL-vHPC | -0.9981 | 0.083333 | 0.862745 | 1.649306 | 4 |
| PL-VTA | 0.484472 | 0.5 | 0.862745 | 76.47569 | 4 |
| IL-aIC | 0.674806 | 0.5 | 0.862745 | 77.17014 | 4 |
| IL-pIC | 0.721221 | 0.25 | 0.862745 | 89.67014 | 4 |
| IL-NAcC | 0.727773 | 0.5 | 0.862745 | 77.17014 | 4 |
| IL-NAcS | -0.83531 | 0.25 | 0.862745 | 10.32986 | 4 |
| IL-BLA | 0.526076 | 0.416667 | 0.862745 | 81.33681 | 4 |
| IL-MD | 0.634098 | 0.416667 | 0.862745 | 81.42361 | 4 |
| IL-LH | -0.2621 | 0.75 | 0.868421 | 34.80903 | 4 |
| IL-vHPC | -0.88526 | 0.166667 | 0.862745 | 6.25 | 4 |
| IL-VTA | -0.00377 | 0.916667 | 0.991803 | 56.33681 | 4 |
| aIC-pIC | 0.31522 | 0.666667 | 0.862745 | 69.01042 | 4 |
| aIC-NAcC | 0.980508 | 0.083333 | 0.862745 | 98.00347 | 4 |
| aIC-NAcS | -0.18164 | 1 | 1 | 47.30903 | 4 |
| aIC-BLA | 0.563656 | 0.5 | 0.862745 | 78.55903 | 4 |
| aIC-MD | -0.08184 | 1 | 1 | 51.30208 | 4 |
| aIC-LH | -0.88876 | 0.083333 | 0.862745 | 1.475694 | 4 |
| aIC-vHPC | -0.81278 | 0.333333 | 0.862745 | 15.10417 | 4 |
| aIC-VTA | 0.578917 | 0.666667 | 0.862745 | 69.53125 | 4 |
| pIC-NAcC | 0.48492 | 0.5 | 0.862745 | 77.95139 | 4 |
| pIC-NAcS | -0.6019 | 0.5 | 0.862745 | 21.26736 | 4 |
| pIC-BLA | 0.837137 | 0.083333 | 0.862745 | 97.56944 | 4 |
| pIC-MD | 0.390935 | 0.583333 | 0.862745 | 73.17708 | 4 |
| pIC-LH | 0.016906 | 0.916667 | 0.991803 | 43.22917 | 4 |
| pIC-vHPC | -0.33434 | 0.666667 | 0.862745 | 31.33681 | 4 |
| pIC-VTA | -0.58764 | 0.416667 | 0.862745 | 19.44444 | 4 |
| NAcC-NAcS | -0.23225 | 1 | 1 | 49.21875 | 4 |
| NAcC-BLA | 0.710609 | 0.333333 | 0.862745 | 85.50347 | 4 |
| NAcC-MD | -0.05735 | 1 | 1 | 52.08333 | 4 |
| NAcC-LH | -0.83532 | 0.083333 | 0.862745 | 2.170139 | 4 |
| NAcC-vHPC | -0.77451 | 0.333333 | 0.862745 | 13.71528 | 4 |
| NAcC-VTA | 0.41207 | 0.666667 | 0.862745 | 68.83681 | 4 |
| NAcS-BLA | -0.14459 | 0.666667 | 0.862745 | 32.55208 | 4 |
| NAcS-MD | -0.95158 | 0.083333 | 0.862745 | 0.954861 | 4 |
| NAcS-LH | -0.28559 | 0.666667 | 0.862745 | 30.55556 | 4 |
| NAcS-vHPC | 0.655404 | 0.416667 | 0.862745 | 82.20486 | 4 |
| NAcS-VTA | 0.297018 | 0.666667 | 0.862745 | 69.61806 | 4 |
| BLA-MD | -0.14053 | 1 | 1 | 48.17708 | 4 |
| BLA-LH | -0.43297 | 0.5 | 0.862745 | 23.00347 | 4 |
| BLA-vHPC | -0.2578 | 0.75 | 0.868421 | 35.41667 | 4 |
| BLA-VTA | -0.27313 | 0.583333 | 0.862745 | 25.78125 | 4 |
| MD-LH | 0.509769 | 0.583333 | 0.862745 | 73.17708 | 4 |
| MD-vHPC | -0.48562 | 0.583333 | 0.862745 | 25.95486 | 4 |
| MD-VTA | -0.33251 | 0.666667 | 0.862745 | 30.46875 | 4 |
| LH-vHPC | 0.503491 | 0.583333 | 0.862745 | 74.56597 | 4 |
| LH-VTA | -0.74188 | 0.333333 | 0.862745 | 14.67014 | 4 |
| vHPC-VTA | -0.44962 | 0.583333 | 0.862745 | 27.77778 | 4 |

Adjusted p-values were obtained from permutation-derived null distributions of Pearson correlation coefficients and corrected for multiple comparisons using the Benjamini–Hochberg false discovery rate (FDR). Percentile values indicate the position of each observed correlation relative to the permutation-based null distribution. *n* indicates the number of animals included in each correlation analysis.

**Table 4.**
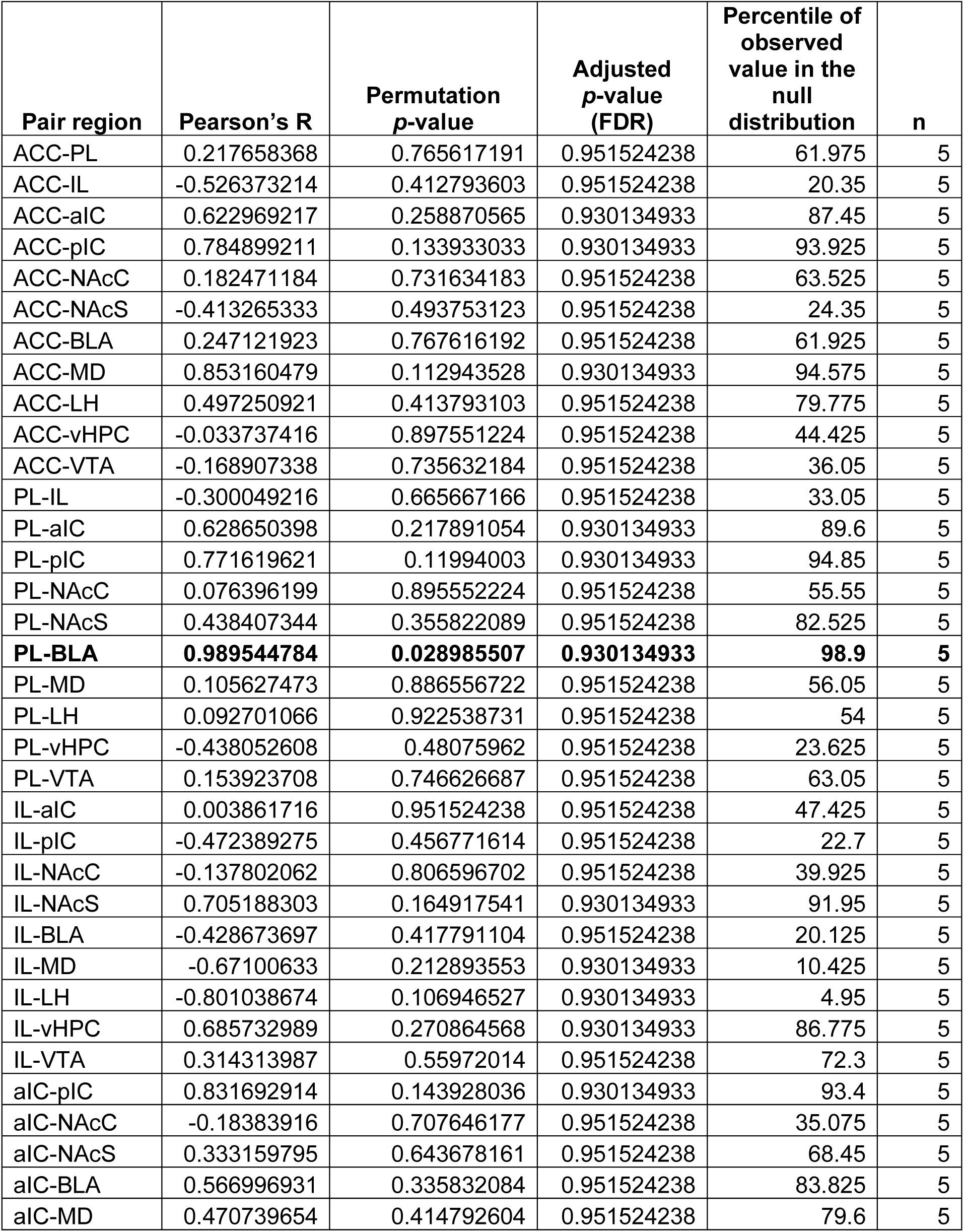
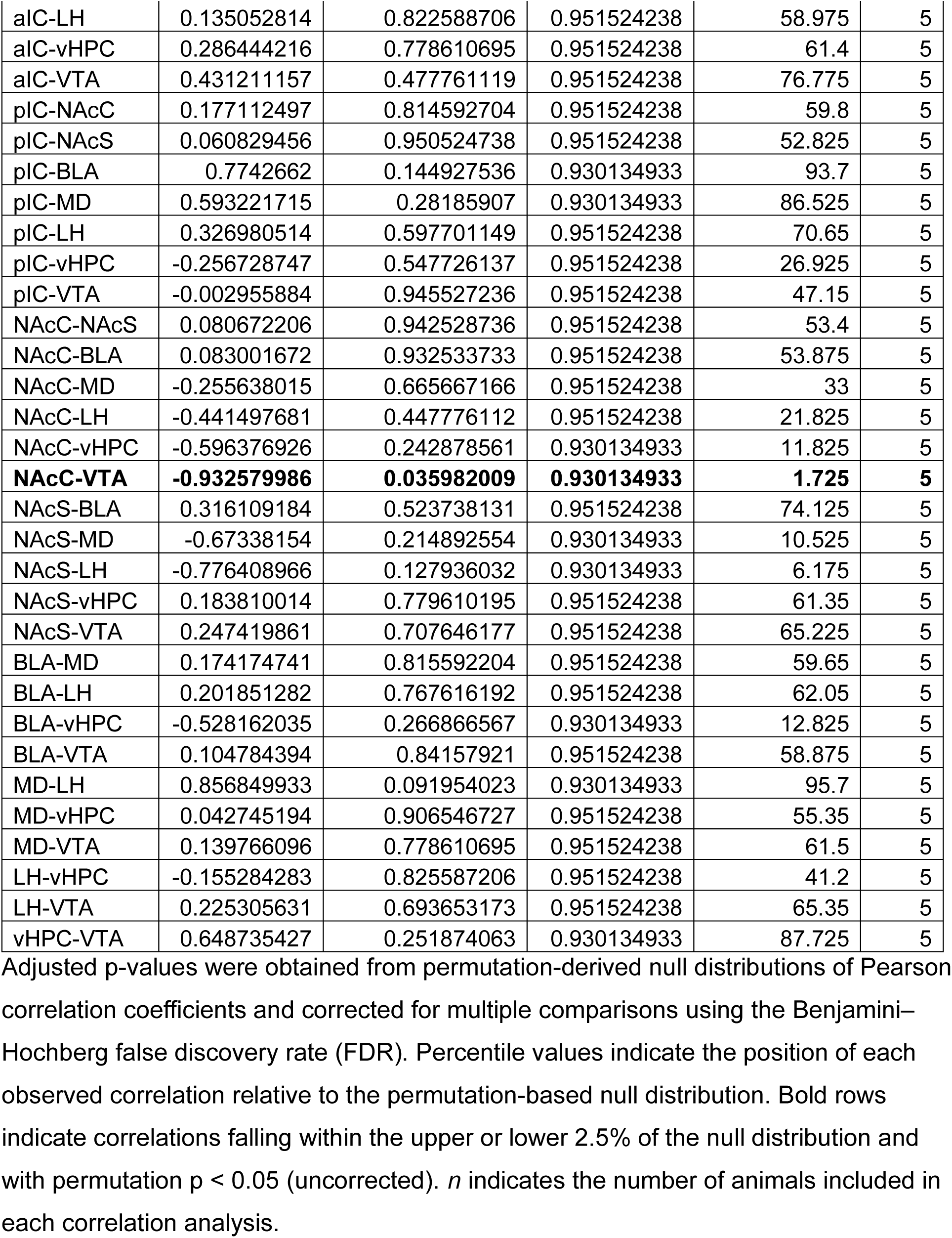
Pearson’s correlations of mean c-Fos + cells count across brain regions during prosocial choices.

**Table 5.**
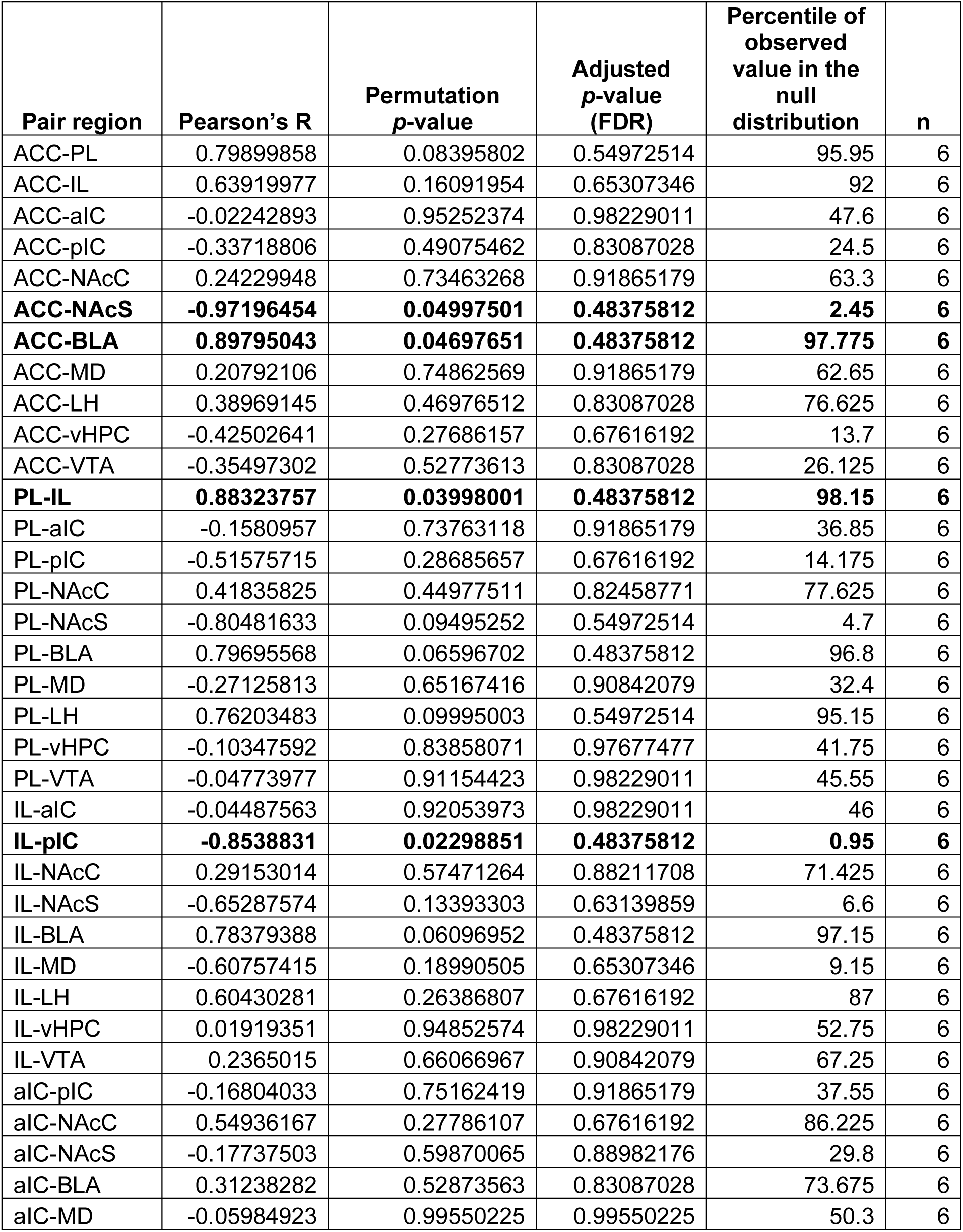
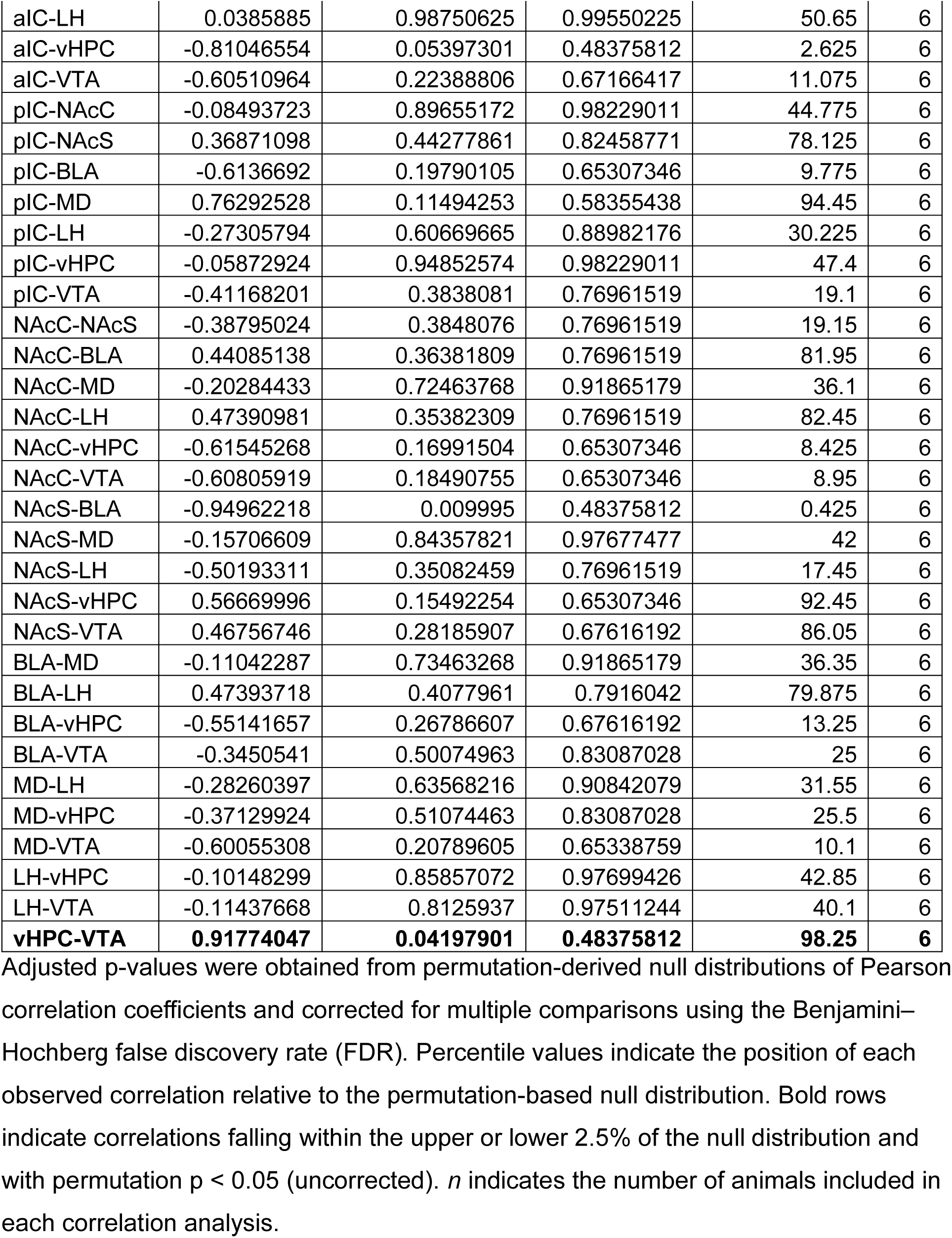
Pearson’s correlations of mean c-Fos + cells count across brain regions during selfish choices.

| Pair region | Pearson's R | Permutation p-value | Adjusted p-value (FDR) | Percentile of observed value in the null distribution | n |
| --- | --- | --- | --- | --- | --- |
| ACC-PL | 0.79899858 | 0.08395802 | 0.54972514 | 95.95 | 6 |
| ACC-IL | 0.63919977 | 0.16091954 | 0.65307346 | 92 | 6 |
| ACC-aIC | -0.02242893 | 0.95252374 | 0.98229011 | 47.6 | 6 |
| ACC-pIC | -0.33718806 | 0.49075462 | 0.83087028 | 24.5 | 6 |
| ACC-NAcC | 0.24229948 | 0.73463268 | 0.91865179 | 63.3 | 6 |
| **ACC-NAcS** | **-0.97196454** | **0.04997501** | **0.48375812** | **2.45** | **6** |
| **ACC-BLA** | **0.89795043** | **0.04697651** | **0.48375812** | **97.775** | **6** |
| ACC-MD | 0.20792106 | 0.74862569 | 0.91865179 | 62.65 | 6 |
| ACC-LH | 0.38969145 | 0.46976512 | 0.83087028 | 76.625 | 6 |
| ACC-vHPC | -0.42502641 | 0.27686157 | 0.67616192 | 13.7 | 6 |
| ACC-VTA | -0.35497302 | 0.52773613 | 0.83087028 | 26.125 | 6 |
| **PL-IL** | **0.88323757** | **0.03998001** | **0.48375812** | **98.15** | **6** |
| PL-aIC | -0.1580957 | 0.73763118 | 0.91865179 | 36.85 | 6 |
| PL-pIC | -0.51575715 | 0.28685657 | 0.67616192 | 14.175 | 6 |
| PL-NAcC | 0.41835825 | 0.44977511 | 0.82458771 | 77.625 | 6 |
| PL-NAcS | -0.80481633 | 0.09495252 | 0.54972514 | 4.7 | 6 |
| PL-BLA | 0.79695568 | 0.06596702 | 0.48375812 | 96.8 | 6 |
| PL-MD | -0.27125813 | 0.65167416 | 0.90842079 | 32.4 | 6 |
| PL-LH | 0.76203483 | 0.09995003 | 0.54972514 | 95.15 | 6 |
| PL-vHPC | -0.10347592 | 0.83858071 | 0.97677477 | 41.75 | 6 |
| PL-VTA | -0.04773977 | 0.91154423 | 0.98229011 | 45.55 | 6 |
| IL-aIC | -0.04487563 | 0.92053973 | 0.98229011 | 46 | 6 |
| **IL-pIC** | **-0.8538831** | **0.02298851** | **0.48375812** | **0.95** | **6** |
| IL-NAcC | 0.29153014 | 0.57471264 | 0.88211708 | 71.425 | 6 |
| IL-NAcS | -0.65287574 | 0.13393303 | 0.63139859 | 6.6 | 6 |
| IL-BLA | 0.78379388 | 0.06096952 | 0.48375812 | 97.15 | 6 |
| IL-MD | -0.60757415 | 0.18990505 | 0.65307346 | 9.15 | 6 |
| IL-LH | 0.60430281 | 0.26386807 | 0.67616192 | 87 | 6 |
| IL-vHPC | 0.01919351 | 0.94852574 | 0.98229011 | 52.75 | 6 |
| IL-VTA | 0.2365015 | 0.66066967 | 0.90842079 | 67.25 | 6 |
| aIC-pIC | -0.16804033 | 0.75162419 | 0.91865179 | 37.55 | 6 |
| aIC-NAcC | 0.54936167 | 0.27786107 | 0.67616192 | 86.225 | 6 |
| aIC-NAcS | -0.17737503 | 0.59870065 | 0.88982176 | 29.8 | 6 |
| aIC-BLA | 0.31238282 | 0.52873563 | 0.83087028 | 73.675 | 6 |
| aIC-MD | -0.05984923 | 0.99550225 | 0.99550225 | 50.3 | 6 |
| aIC-LH | 0.0385885 | 0.98750625 | 0.99550225 | 50.65 | 6 |
| aIC-vHPC | -0.81046554 | 0.05397301 | 0.48375812 | 2.625 | 6 |
| aIC-VTA | -0.60510964 | 0.22388806 | 0.67166417 | 11.075 | 6 |
| pIC-NAcC | -0.08493723 | 0.89655172 | 0.98229011 | 44.775 | 6 |
| pIC-NAcS | 0.36871098 | 0.44277861 | 0.82458771 | 78.125 | 6 |
| pIC-BLA | -0.6136692 | 0.19790105 | 0.65307346 | 9.775 | 6 |
| pIC-MD | 0.76292528 | 0.11494253 | 0.58355438 | 94.45 | 6 |
| pIC-LH | -0.27305794 | 0.60669665 | 0.88982176 | 30.225 | 6 |
| pIC-vHPC | -0.05872924 | 0.94852574 | 0.98229011 | 47.4 | 6 |
| pIC-VTA | -0.41168201 | 0.3838081 | 0.76961519 | 19.1 | 6 |
| NAcC-NAcS | -0.38795024 | 0.3848076 | 0.76961519 | 19.15 | 6 |
| NAcC-BLA | 0.44085138 | 0.36381809 | 0.76961519 | 81.95 | 6 |
| NAcC-MD | -0.20284433 | 0.72463768 | 0.91865179 | 36.1 | 6 |
| NAcC-LH | 0.47390981 | 0.35382309 | 0.76961519 | 82.45 | 6 |
| NAcC-vHPC | -0.61545268 | 0.16991504 | 0.65307346 | 8.425 | 6 |
| NAcC-VTA | -0.60805919 | 0.18490755 | 0.65307346 | 8.95 | 6 |
| NAcS-BLA | -0.94962218 | 0.009995 | 0.48375812 | 0.425 | 6 |
| NAcS-MD | -0.15706609 | 0.84357821 | 0.97677477 | 42 | 6 |
| NAcS-LH | -0.50193311 | 0.35082459 | 0.76961519 | 17.45 | 6 |
| NAcS-vHPC | 0.56669996 | 0.15492254 | 0.65307346 | 92.45 | 6 |
| NAcS-VTA | 0.46756746 | 0.28185907 | 0.67616192 | 86.05 | 6 |
| BLA-MD | -0.11042287 | 0.73463268 | 0.91865179 | 36.35 | 6 |
| BLA-LH | 0.47393718 | 0.4077961 | 0.7916042 | 79.875 | 6 |
| BLA-vHPC | -0.55141657 | 0.26786607 | 0.67616192 | 13.25 | 6 |
| BLA-VTA | -0.3450541 | 0.50074963 | 0.83087028 | 25 | 6 |
| MD-LH | -0.28260397 | 0.63568216 | 0.90842079 | 31.55 | 6 |
| MD-vHPC | -0.37129924 | 0.51074463 | 0.83087028 | 25.5 | 6 |
| MD-VTA | -0.60055308 | 0.20789605 | 0.65338759 | 10.1 | 6 |
| LH-vHPC | -0.10148299 | 0.85857072 | 0.97699426 | 42.85 | 6 |
| LH-VTA | -0.11437668 | 0.8125937 | 0.97511244 | 40.1 | 6 |
| **vHPC-VTA** | **0.91774047** | **0.04197901** | **0.48375812** | **98.25** | **6** |

Adjusted p-values were obtained from permutation-derived null distributions of Pearson correlation coefficients and corrected for multiple comparisons using the Benjamini–Hochberg false discovery rate (FDR). Percentile values indicate the position of each observed correlation relative to the permutation-based null distribution. Bold rows indicate correlations falling within the upper or lower 2.5% of the null distribution and with permutation $p < 0.05$ (uncorrected). $n$ indicates the number of animals included in each correlation analysis.

## DISCUSSION

Here, we reveal strain-dependent differences in social decision-making: CD1 mice favored selfish choices, whereas C57BL/6 mice were more prosocial. CD1 mice showed higher mPFC and NAcC c-Fos expression, with mPFC activity correlating with selfish behavior. Network analyses identified coordinated PL–IL–NAcC–VTA engagement during social decision-making, indicating that distinct social strategies recruit different prefrontal–striatal–midbrain circuits.

We adapted a validated social decision-making task (Scheggia et al., 2022) by adding Pavlovian cue conditioning and forced sampling to ensure balanced option experience before free choice. This design aligns with established decision-making frameworks (St Onge and Floresco, 2009; Simon et al., 2009) and facilitates future electrophysiological studies through cue-aligned analyses. Combined with c-Fos mapping, it enabled assessment of distributed cortical–subcortical activity underlying socially biased decisions.

### Selfish and prosocial behavior are strain-dependent

Recent studies have established mice as a robust model for investigating prosocial behaviors (Michael J.M. Gachomba et al. 2024). In particular, C57BL/6 mice display robust prosocial behaviors, including consolation and affiliative behaviors toward distressed conspecifics (Wu et al., 2021; Zhang et al., 2024), helping trapped cagemates (Pozo et al., 2023), avoiding harm to others (Song et al., 2023), and sharing rewards with partners (Scheggia et al., 2022; Misiołek et al., 2023). Consistent with these findings, dominant C57BL/6 mice displayed a robust prosocial phenotype in our modified social decision-making paradigm. In contrast, CD1 mice exhibited a pronounced bias toward selfish choices.

Several factors may contribute to this divergence. First, social interaction is a key determinant of prosocial expression in mice (Scheggia et al. 2022; Song et al. 2023). Here, C57BL/6 mice engaged in more social interaction with their partners than CD1 mice. This further supports strain-differences in baseline pro-sociability, as CD1 mice interacted less and were less likely to provide rewards to partner mice. Second, CD1 mice display higher levels of agonistic behaviors, including aggressive interactions, than other strains (Hsieh et al. 2017; Van Loo et al. 2003; Cum et al. 2024). Agonistic behaviors are indicators of social dominance and hierarchical status in mice (Zhou et al. 2018; Fulenwider et al. 2022). Consistent with this framework, CD1 mice exhibit elevated dominance-associated behaviors during social competition and increased territorial marking compared to C57BL/6 mice (Cum et al., 2024). Given that dominant mice were used as subjects in social decision-making, the selfish bias observed in CD1 mice may reflect dominance-related tendencies that prioritize individual reward even in the absence of competition. Future studies should determine whether subordinate CD1 mice display different social decision-making profiles. Finally, motivational differences may further bias social choices. CD1 mice exerted greater effort to obtain rewards than C57BL/6 mice (Campos-Ordoñez and Buriticá 2024), indicating heightened reward sensitivity. This greater valuation of individual benefits could amplify competitive tendencies, fostering selfish decisions in CD1 mice.

Taken together, these findings indicate that selfish and prosocial behaviors are not uniform features of murine social decision-making but instead vary across strains. While dominance or motivational traits were not directly evaluated here, strain-dependent behavioral tendencies reported in the literature may explain these differences during social decision-making. Such behavioral divergence provides a useful framework for interpreting the strain-specific neural activity patterns observed across cortical–subcortical circuits.

### Selective mPFC engagement underlies selfish choice bias

The mPFC is a hub for integrating decision-making (Gangopadhyay et al. 2021), and its subregions, such as ACC and PL, have been implicated in prosocial behavior (Song et al. 2023; Conde-Moro et al. 2024). Our data, however, show that CD1 mice exhibit higher c-Fos activation in these mPFC subregions than C57BL/6 mice, and that mPFC activity is inversely correlated with prosocial choice. Thus, mPFC engagement appears preferentially linked to selfish decisions, contrary to previous reports. However, notably, the vast majority of prosocial studies in mice use C57BL/6 mice; thus, findings may unintentionally be biased towards phenotypically prosocial breeds.

In humans, the medial prefrontal cortex (mPFC) plays a central role in integrating self-interest during social decision-making. Functional neuroimaging studies demonstrate that mPFC activity increases with the level of self-risk during altruistic choice, indicating enhanced processing of self-related costs when individuals consider acting in their own interest (Hu et al. 2017). These findings suggest that mPFC does not exclusively promote prosocial behavior but rather encodes decision variables related to self-interest within a social context. Importantly, self-interest in social contexts often unfolds within hierarchical environments, where individuals must balance personal gain against social consequences (Sapolsky 2005). Consistent with this broader role, medial prefrontal regions that are functionally conserved across species, including rodent ACC, PL, and IL, are critically involved in regulating social hierarchy and dominance-related behaviors (Wang et al. 2011). Activity within these subregions in rodents influences competitive interactions and hierarchical status. For instance, optogenetic activation of PL drives competitive dominance (Zhou et al. 2018), whereas ACC and PL activity are implicated when mice successfully compete for rewards (Li et al. 2022; Padilla-Coreano et al. 2022), highlighting mPFC contributions to social dominance in different competitive contexts. Given that our subject mice were of dominant rank, the elevated c-Fos expression observed in ACC, PL, and IL during selfish choice in CD1 mice may reflect recruitment of medial prefrontal circuits integrating self-interest with competitive social interactions. Together, our findings suggest that increased mPFC activation may reflect the implementation of a dominance-aligned, resource-maximizing strategy within a social context.

Although IC is implicated in prosocial behavior (Cox et al., 2022; Rogers-Carter et al., 2018), we found no significant IC recruitment during either prosocial or selfish choices. IC activity is associated with emotional contagion and affective processing in social contexts of distress (Rogers-Carter and Christianson, 2019; Keysers et al., 2022). Because our task did not present overt vulnerability or distress, the IC may have remained largely unengaged. Overall, these results highlight that strain-dependent social strategies are linked to engagement of the mPFC during social decision-making.

### NAcC is engaged in CD1 selfish mice more than C57BL/6 prosocial mice

The nucleus accumbens (NAc) has been related to social behaviors including cooperative interactions (Ben-Ami Bartal et al. 2021; Conde-Moro et al. 2024; Hazani et al. 2025). In our study, however, c-Fos expression was markedly higher in the NAcC of selfish CD1 mice than in C57BL/6 mice, indicating that this region is more engaged during self-oriented decision-making. This strain-dependent activity pattern may be related to previously described differences in social dominance traits. Prior work shows that inhibiting the NAc (core and shell) reduces social dominance (Hollis et al. 2015; Shan et al. 2022), indicating its role in competitive behavior. Accordingly, elevated NAcC activity during selfish choices in CD1 mice may reflect striatal engagement in competitive reward acquisition rather than prosocial valuation.

Strain differences were not observed across the other subcortical regions evaluated, NAc shell, BLA, MD, LH, vHPC, and VTA. Although these areas have been implicated in prosocial and dominance behaviors (Hung et al. 2017; Phillips et al. 2019; Scheggia et al. 2022; Padilla-Coreano et al. 2022; Song et al. 2023), we detected no significant c-Fos differences during social choice. Rather than ruling out their role, this suggests that social decisions depend more on coordinated circuit interactions than on isolated regional activation. Indeed, the absence of strain-level differences in these regions is consistent with their participation in the broader social decision-making network identified here, where their functional relevance may be expressed through coordinated activity with other regions rather than through differential recruitment by strain.

### Network-Level Organization of Social Decision-Making Circuits

Network-level analysis revealed coordinated activity across mPFC, striatal, and midbrain regions. Rather than reflecting isolated subregional effects, social decision-making was associated with subnetwork interactions among PL, IL, NAcC, and VTA. Although individual pairwise correlations did not survive multiple-comparison correction, the correlation values are strong, and a permutation test supports functionally meaningful coupling across these regions. This pattern is consistent with extensive evidence in rodents showing that cortico-striatal and mesolimbic circuits coordinate reward-based and goal-directed decision-making processes (Lammel et al. 2014; Floresco 2015; Khani and Rainer 2016). Importantly, cortico-striatal and mesolimbic circuits have been shown to regulate socially motivated behaviors, including prosocial and competitive interaction actions (Choi et al., 2024; Conde-Moro et al., 2024; Xing et al., 2021), suggesting that social choices may emerge from the recruitment of canonical value-based decision networks within a social context.

Network-based statistical analyses and permutation tests of pairwise interactions suggest that prosocial, selfish, and no-preference mice exhibit distinct patterns of subnetwork interactions. Prosocial mice exhibited coordinated PL–BLA activity patterns consistent with previous findings (Scheggia et al. 2022). Notably, we observed a negative correlation between VTA and NAcC. While this finding does not establish directionality or mechanism, it raises the possibility that social decision-making involves dynamic regulatory interactions within this circuit, potentially reflecting VTA GABAergic inhibition over NAc during reward-seeking in stress conditions (Lowes et al., 2021). In selfish-preferring mice, negative associations between ACC–NAcS and NAcS–BLA emerged as part of a significant coordinated subnetwork. Although ACC-NAcS has not yet been explored in social behaviors, ACC to NAc projections have been implicated in social transfer of pain (Smith et al. 2021), a form of emotional contagion essential for prosocial behavior in rodents (Scheggia et al. 2022). Within this framework, the negative association between ACC and NAcS observed may reflect reduced coordination within circuits supporting emotional contagion during selfish choices. Similarly, a prior work showed increased BLA–NAc coherence during cooperative behavior (Conde-Moro et al. 2024), whereas we observed an inverse pattern in selfish mice, suggesting distinct modes of amygdala–striatal engagement during social decisions. Finally, no-preference mice showed no significant pairwise correlations nor subnetwork activity, suggesting that coordinated network activity is specifically associated with the expression of social decision preferences.

### Limitations of the study and future directions

Several limitations should be considered. First, c-Fos mapping provides indirect, temporally limited measures of neuronal activation, thereby precluding conclusions about synaptic directionality or causal interactions. Second, phenotypic subgroup analyses were conducted with relatively small sample sizes, and these associations lay the groundwork for further investigation to confirm network states based on social choices. Future experiments combining circuit manipulations and in vivo multisite electrophysiological recordings will be necessary to determine the causal contributions and temporal dynamics of the identified corticostriatal–midbrain circuits during social decision-making. Third, the present study was conducted exclusively in male mice, given that past reports indicate that in similar operant tasks, female mice are less prosocial (Scheggia et al. 2022). Future studies including female subjects will be important for determining the generalizability of these findings.

## CONCLUSION

Our findings reveal strain-dependent differences in social decision-making, with CD1 mice exhibiting greater selfish behavior, accompanied by elevated mPFC and NAcC activity. At the network level, social decision-making engaged coordinated VTA–NAc–prefrontal interactions, while prosocial and selfish mice recruited distinct cortical–subcortical configurations. Together, these results provide evidence that social decision-making is supported by distributed cortical and subcortical network dynamics that differentially shape selfish and prosocial choices.

## METHODS

### Animals

All experimental animals were maintained on a 12 h reverse light cycle (lights off: 8:30 A.M.–8:30 P.M.), and all behavioral testing was conducted during the dark phase between 9:00 A.M. and 6:00 P.M. Male CD1 (n = 24; Charles River Laboratories) and C57BL/6 (n = 24; The Jackson Laboratory) mice were 8 weeks old upon arrival and group-housed (4 per cage). Behavioral experiments began at least one week after arrival. Mice were handled for at least 2 days prior to testing. Throughout the experiment, animals were food-restricted to 90% of their baseline body weight, with water available ad libitum. All procedures were performed in accordance with the University of Florida regulations and were approved by the Institutional Animal Care and Use Committee (IACUC).

### Behavioral assays

#### Social decision-making task

C57BL/6 and CD1 mice (16 mice per strain) were trained individually in Med-PC operant chambers connected to a custom 3D-printed triangular chamber. Mice placed in the operant chambers were designated as subjects, whereas their cagemates were positioned in the adjacent triangular compartment as partners (**Figure 1 A**). The two compartments were separated by a metal mesh (1 cm openings) that allowed direct social interaction. Each chamber was equipped with a modified 3D-printed reward port connected to a syringe pump (Med Associates, PHM-210), which delivered 15 μl of vanilla Ensure per reward. Each session lasted up to 40 minutes and was controlled using custom-written Med-PC programs. Operant chambers and partners’ compartments were cleaned using soap, water, and 70% of alcohol between animals.

#### Social decision-making training

All mice underwent training for the social decision-making task, which consisted of three consecutive stages: 1) Reward conditioning, 2) Forced-choice training, and 3) Free-choice training.

1. Reward conditioning **(Figure 1C**): Subjects and partners were confined to their respective compartments and trained to associate two distinct auditory cues, serving as conditioned stimuli (CS), with prosocial (reward delivered to both mice) or selfish (reward delivered to the subject only) outcomes. Tone–outcome pairings were fully counterbalanced across mice, with contingencies reversed between animals to prevent cue bias. The CS–outcome assignments established during this phase were maintained throughout subsequent training stages. Each session consisted of 30 trials of a single CS–outcome contingency (either prosocial or selfish). In each trial, the CS was presented for 10 s, and reward delivery occurred 5 s after CS onset, midway through cue presentation. Trials were separated by a variable inter-trial interval (ITI) of 1–3 min. Contingencies alternated across 6 consecutive days, with 3 days of exclusive exposure to each outcome (prosocial: days 1, 3, and 5; selfish: days 2, 4, and 6). Learning performance was assessed by quantifying the number of reward port entries during CS presentation.
2. Forced-choice training (**Figure 1F**): Subjects and partners were confined to their respective compartments, with the subject having access to two nose-poke ports. Mice learned to associate prosocial and selfish outcomes with their respective CSs. The assignment of prosocial and selfish contingencies to the left or right nose-poke was counterbalanced across mice and remained fixed for each animal throughout subsequent sessions and training stages. In each session, only one nose-poke was designated as active, whereas responses at the other port had no programmed consequences. The active nose-poke was illuminated to indicate availability, while the inactive port remained unlit. Responses at the active nose-poke triggered presentation of the associated CS (5 s duration), followed by reward delivery to both mice (prosocial) or to the subject only (selfish). The reward was delivered 2.5 s after CS onset, midway through cue presentation. Following each reinforced response, a 5-s timeout period was imposed, during which additional nose-poke responses did not trigger CS presentation or reward delivery. Prosocial and selfish sessions were alternated across 6 consecutive days, with 3 days of exclusive exposure to each outcome (prosocial: days 1, 3, and 5; selfish: days 2, 4, and 6). Learning performance was assessed by quantifying responses at active and inactive nose-pokes across trials.
3. Free-choice trials (**Figure 1I**): Subjects and partners were confined to their respective compartments and first received a block of forced trials, followed by 30 min of free-choice trials during which both nose-pokes were available (indicated by illumination of both ports). Forced trials were conducted as described above and terminated once each mouse had received 20 rewards for each contingency (prosocial and selfish). The order of prosocial and selfish forced-trial blocks was counterbalanced across days and across animals. At the conclusion of the forced block, a 30 s inter-trial interval (ITI) preceded the onset of free-choice trials. During free-choice trials, responses at either nose-poke triggered presentation of the corresponding CS (prosocial or selfish) and reward delivery as previously described. The 5-s post-reinforcement timeout described above was maintained throughout this phase. Mice were trained for 8 consecutive days until they exhibited a stable preference for one contingency for at least 3 consecutive days. Learning performance was assessed by quantifying responses at the prosocial or selfish nose-pokes.

#### Alone decision-making task

C57BL/6 and CD1 mice in the alone condition were trained with the same apparatus and trial structure as in the social decision-making task, except that no partner was present (Figure 1B). Mice in this condition were naïve to the social setup and had no prior experience with the prosocial contingency. To maintain consistency with the social condition, nose-pokes retained their original CS–side assignments and were analyzed as Option 1 and Option 2. These labels referred solely to the two counterbalanced stimulus–side configurations established during training. Specifically, C1 was assigned so that half of the animals experienced the conditioned stimulus (CS) on the left nose-poke and half on the right, whereas C2 represented the complementary counterbalanced configuration. Thus, in the alone condition, Option 1 and Option 2 did not reflect social preference but were defined exclusively by stimulus identity and side assignment.

#### Social interactions

Social interaction bouts, defined as any face-to-face encounter between the subject and recipient through the metal mesh barrier lasting at least 1 second, were quantified during the c-Fos test using manual scoring in BORIS software by an experimenter blind to strain and experimental condition. Behaviors such as proximity without direct face orientation were not included. The total number of social interaction bouts was quantified across the entire session.

#### Social rank assays

Given that prior studies have demonstrated that social rank influences social decision-making, (Gachomba et al. 2022; Scheggia et al. 2022), we controlled for this variable by measuring social ranks and using dominant mice as subjects and subordinate mice as partners in both mouse strains. To assess social rank, we used the tube test for C57BL/6 mice and the urine-marking assay for CD1 mice, based on prior evidence that these tests reliably measure dominance hierarchies in their respective strains (Cum et al. 2024).

Tube test: All training and testing were conducted in a dimly lit (20 lux), quiet room. Mice underwent 3 consecutive days of tube training prior to testing. During training, mice were guided to traverse the tube, turn at the opposite end, and return to the starting point, completing ten full crossings per session. A plastic cylinder slightly smaller than the tube was used, when necessary, to gently prevent retreat or encourage forward movement. During testing, each mouse was paired once per session with each of its cage mates. The loser was defined as the first animal to exit the tube with all four paws, whereas the winner was defined as the animal that remained inside the tube. The order of pairings and starting positions was counterbalanced across days. Tube testing was conducted for at least 8 consecutive days and continued until stable rank hierarchies were maintained for at least 3 consecutive days.

Urine marking test: CD1 mice were tested in a dimly lit room (20 lux) within a clear plastic enclosure (16.5 cm × 39.0 cm) without flooring. The enclosure was divided into two equal compartments by a clear, perforated partition that allowed visual and olfactory interaction. Chromatography paper (Whatman cellulose) was placed beneath the enclosure and covered with a metal mesh to prevent chewing. Mice were tested against different cage mates across 6 consecutive days, with sessions lasting 3 h each. Urine markings were quantified under UV illumination by trained observers. Each pair within a cage was tested in 1–2 sessions, and each mouse was tested no more than once per day. The number of urine marks per session was used to determine territorial-based social rank. Matches were considered tied when the difference between animals was <20% or when there were fewer than 5 urine spots.

#### Food restriction control group

A separate cohort of C57BL/6 and CD1 mice was food-restricted for 20 days and subsequently perfused to serve as a baseline c-Fos control group. Mice were maintained at 90% of their initial body weight and were provided with 3 g of standard chow daily at a consistent time each day. Animals were handled for at least 5 consecutive days prior to perfusion.

### Histology

On the last day of training, subject mice (C57BL/6 mice: social, n=7, alone n=4; CD1 mice: social, n=8, alone n= 4) or food restricted control mice (n=8 per strain) were perfused 90 minutes after the session via intraperitoneal injection of pentobarbital (31.5 mg/ml), followed by transcardial perfusion with 0.9% sodium chloride solution and 4% paraformaldehyde (PFA) in 1x phosphate-buffered saline (PBS). Brains were extracted and post-fixed in 4% PFA for 24 hours and then cryoprotected in 30% sucrose in 1x PBS until the brain sank to the bottom of the tube. Brains were then sliced at 40 μm using a microtome, and slices were stored at 4°C in 0.05% sodium azide until staining.

### Immunohistochemistry

Brain sections were incubated for 1 h in a blocking solution consisting of PBS supplemented with 3% normal donkey serum (NDS) and 0.2% Triton X-100. Sections were then incubated overnight at 4°C with a rabbit monoclonal recombinant IgG primary antibody (Synaptic Systems; 1:5,000 dilution in PBS-T containing 3% NDS). The next day, sections were incubated for 2 hours at room temperature in goat anti-rabbit IgG (H+L) cross-absorbed secondary antibody conjugated with Alexa Fluor 555 (Thermo Fisher Scientific; dilution 1:500 in PBS-T with 3% NDS). After washing, sections were mounted using Fluoromount-G with DAPI (Thermo Fisher Scientific). For each brain region, three sections were collected at anterior, medial, and posterior coordinates, with the sampled hemisphere counterbalanced across animals. Regions of interest were manually delineated and imaged at 20× magnification using a fluorescence microscope, ensuring consistent sampling of representative areas. Imaging parameters were kept constant across all animals and regions. c-Fos–positive cells were quantified for each section, and the mean count per region was calculated by averaging across the three sections.

### Data collection and analysis

#### Social and alone decision-making task

Learning performance was evaluated at each training stage. For stage1, the probability of the reward port was calculated as the proportion of trials in which at least one port entry occurred within the 10 s window following CS onset (0–10 s), defined as Probability entry= (Number of trials with ≥1 entry)/ (Total number of trials). Reward port entry latency was defined as the time to the first port entry minus CS onset within the same 10 s window (0–10 s). Mean latency was calculated by averaging across trials with at least one entry for each subject and day. For stages 2 and 3, the number of nose-poke responses was quantified using MED-PC software (Med Associates). To assess discrimination between active and inactive nose-pokes, a learning index was calculated as: Index learning = (Number of active nose-pokes − Number of inactive nose-pokes) / (Total number of nose-pokes), where values close to 1 indicate preferential responding at the active nose-poke, values close to −1 indicate preferential responding at the inactive nose-poke, and values near 0 indicate no discrimination. To quantify individual preferences for prosocial over selfish responses, a decision preference score was calculated as: Preference score= (number of prosocial responses – number of selfish responses)/ (total number of responses), wherein values close to 1 indicate prosocial preference, values close to 0 indicate no-preference, and values close to -1 indicate selfish preference. To assess side and tone preferences, the proportion of prosocial and selfish choices was calculated as: Percentage choice = (Number of prosocial or selfish responses × 100) / (Total number of responses).

#### Immunoreactivity quantification

Brain sections were imaged at 20× magnification using a KEYENCE BZ-X800 all-in-one fluorescence microscope (Keyence Corporation of America) under high-sensitivity settings. Maximum projection images were generated using the Full Focus option. Images were generated for the ACC (bregma 1.53–0.85 mm), PL (bregma 2.09–1.41 mm), IL (bregma 1.97–1.41 mm), aIC (bregma 1.53–0.85 mm), pIC (bregma 0.73–0.47 mm), NAcC (bregma 1.53–0.97 mm), NAcS (bregma 1.69–0.73 mm) MD (bregma −0.59 to −1.79 mm), BLA (bregma −1.07 to −1.79 mm), LH (bregma −0.71 to −1.79 mm), vHPC(bregma −2.79 to −3.51 mm), and VTA (-3.15 to -3.79 mm; Paxinos and Franklin’s The Mouse Brain, 5th edition, 2019). c-Fos^+^ cells were hand-counted using the ImageJ Macro Cell Counter plugin. Positive Fos-like immunoreactivity was indicated by strong cell nuclei staining. Cell counts were performed by two experimenters blinded to experimental condition, with region and condition assignments counterbalanced across raters. For each animal, counts were averaged across anterior, medial, and posterior rostrocaudal levels to obtain a single mean c-Fos+ value per brain region.

#### Statistical analysis

Statistical analyses included Shapiro–Wilk tests for normality, Pearson’s correlations, and linear regression lines were used to display best-fit trends in scatterplots. Cell counts across brain regions were correlated with the percentage of prosocial choices (Figure 2–3), number of rewards consumed (Figure S3), and number of social interactions (Figure S4), using Pearson’s test. For these behavioral correlations, p values were adjusted using the Benjamini–Hochberg false discovery rate (FDR) procedure. For network analyses, we used a permutation test (scipy.stats.permutation_test). For each correlation, we generated a null distribution by randomly shuffling cell densities within regions across 2,000 permutations while calculating Pearson’s correlation coefficients. For each permutation, subject labels were shuffled within a given brain region, thus maintaining the mean c-Fos values per region. The Benjamini–Hochberg FDR procedure was applied to two-tailed p values derived from the permutation test. For cross-regional correlations, FDR adjustments were applied to all brain region pairs within each experimental context separately (social decision-making, selfish, prosocial, no-preference, and alone). Although given small sample sizes no individual region pairs survived FDR correction (q < 0.05), correlations within the top and bottom 2.5% percentile of the null distribution and had p<0.05 in the permutation test were reported in network graphs to visualize strong cross-regional associations that were not random based on the permutation test.

To assess coordinated network-level organization, we used a component-based permutation procedure inspired by the Network-Based Statistic (NBS) framework (Zalesky et al. 2010). Pairwise Pearson correlations were computed across all selected brain regions (12 regions; 66 unique region pairs). A primary threshold of p < 0.01 (uncorrected) was used to define suprathreshold edges, and connected components were identified based on edge extent (number of edges). Component significance was assessed using 5,000 permutations. For each permutation, animal values were independently shuffled within each brain region (column-wise permutation, preserving missing values), thereby disrupting cross-regional dependencies while preserving the marginal distributions of each region. For each permuted dataset, the correlation matrix was recomputed and the maximal component extent recorded to generate a null distribution. The component-level p-value (p_component) was defined as the proportion of permutations yielding a component extent greater than or equal to the observed extent (with a +1 correction: (k + 1)/(n_perm + 1). Component-based permutation tests were performed separately within each condition/subgroup to assess the presence of coordinated network organization.

## Author contributions

E.I-H and N.P-C designed research; E.I-H and A.V performed the experiments; E.I-H and M.C analyzed the data; E.I-H and N.P-C wrote the paper.

## Acknowledgments

This research was supported by R01MH139895, McKnight Neuroscience Award, and Klingenstein-Simons Neuroscience Award to N.P-C. We thank Genesis De Leon Rivera, Logan Taylor, and Zishu Zhao (Catherine) for technical assistance.

## References

Ben-Ami Bartal, Inbal, Jocelyn M. Breton, Huanjie Sheng, et al. 2021. “Neural Correlates of Ingroup Bias for Prosociality in Rats.” eLife 10 (July): e65582. 10.7554/eLife.65582.

Burgos-Robles, Anthony, Hector Bravo-Rivera, and Gregory J. Quirk. 2013. “Prelimbic and Infralimbic Neurons Signal Distinct Aspects of Appetitive Instrumental Behavior.” PLoS ONE 8 (2): e57575. 10.1371/journal.pone.0057575.

Campos-Ordoñez, Tania, and Jonathan Buriticá. 2024. “Assessment of the Inbred C57BL/6 and Outbred CD1 Mouse Strains Using a Progressive Ratio Schedule during Development.” Physiology & Behavior 277 (April): 114485. 10.1016/j.physbeh.2024.114485.

Conde-Moro, A. R., F. Rocha-Almeida, E. Gebara, J. M. Delgado-García, C. Sandi, and A. Gruart. 2024. “Involvement of Prelimbic Cortex Neurons and Related Circuits in the Acquisition of a Cooperative Learning by Pairs of Rats.” Cognitive Neurodynamics 18 (5): 2637–58. 10.1007/s11571-024-10107-y.

Cox, Stewart S., Angela M. Kearns, Samuel K. Woods, Brogan J. Brown, Samantha J. Brown, and Carmela M. Reichel. 2022. “The Role of the Anterior Insula during Targeted Helping Behavior in Male Rats.” Scientific Reports 12 (1): 3315. 10.1038/s41598-022-07365-3.

Cum, Meghan, Jocelyn A. Santiago Pérez, Ryo L. Iwata, et al. 2024. “A Multiparadigm Approach to Characterize Dominance Behaviors in CD1 and C57BL6 Male Mice.” Eneuro 11 (11): ENEURO.0342-24.2024. 10.1523/ENEURO.0342-24.2024.

Dai, Bing, Fangmiao Sun, Xiaoyu Tong, et al. 2022. “Responses and Functions of Dopamine in Nucleus Accumbens Core during Social Behaviors.” Cell Reports 40 (8): 111246. 10.1016/j.celrep.2022.111246.

De Paula Cunha Almeida, Caroline, Meghan Cum, Elizabeth Illescas-Huerta, et al. 2025. “Prefrontal and Subcortical C-Fos Mapping of Reward Responses across Competitive and Social Contexts.” Eneuro 12 (11): ENEURO.0158-25.2025. 10.1523/ENEURO.0158-25.2025.

Floresco, Stan B. 2015. “The Nucleus Accumbens: An Interface Between Cognition, Emotion, and Action.” Annual Review of Psychology 66 (1): 25–52. 10.1146/annurev-psych-010213-115159.

Fulenwider, Hannah D., Maya A. Caruso, and Andrey E. Ryabinin. 2022. “Manifestations of Domination: Assessments of Social Dominance in Rodents.” *Genes*, Brain and Behavior 21 (3): e12731. 10.1111/gbb.12731.

Gachomba, Michael J.M., Joan Esteve-Agraz, and Cristina Márquez. 2024. “Prosocial Behaviors in Rodents.” Neuroscience & Biobehavioral Reviews 163 (August): 105776. 10.1016/j.neubiorev.2024.105776.

Gachomba, Michael Joe Munyua, Joan Esteve-Agraz, Kevin Caref, et al. 2022. “Multimodal Cues Displayed by Submissive Rats Promote Prosocial Choices by Dominants.” Current Biology 32 (15): 3288–3301.e8. 10.1016/j.cub.2022.06.026.

Gangopadhyay, Prabaha, Megha Chawla, Olga Dal Monte, and Steve W. C. Chang. 2021. “Prefrontal–Amygdala Circuits in Social Decision-Making.” Nature Neuroscience 24 (1): 5–18. 10.1038/s41593-020-00738-9.

Hazani, Reut, Jocelyn M. Breton, Estherina Trachtenberg, et al. 2025. “Neural and Behavioral Correlates of Individual Variability in Rat Helping Behavior: A Role for Social Affiliation and Oxytocin Receptors.” The Journal of Neuroscience 45 (22): e0845242025. 10.1523/JNEUROSCI.0845-24.2025.

Hollis, Fiona, Michael A. Van Der Kooij, Olivia Zanoletti, Laura Lozano, Carles Cantó, and Carmen Sandi. 2015. “Mitochondrial Function in the Brain Links Anxiety with Social Subordination.” Proceedings of the National Academy of Sciences 112 (50): 15486–91. 10.1073/pnas.1512653112.

Hsieh, Lawrence S., John H. Wen, Laura Miyares, Paul J. Lombroso, and Angélique Bordey. 2017. “Outbred CD1 Mice Are as Suitable as Inbred C57BL/6J Mice in Performing Social Tasks.” Neuroscience Letters 637 (January): 142–47. 10.1016/j.neulet.2016.11.035.

Hu, Jie, Yue Li, Yunlu Yin, Philip R. Blue, Hongbo Yu, and Xiaolin Zhou. 2017. “How Do Self-Interest and Other-Need Interact in the Brain to Determine Altruistic Behavior?” NeuroImage 157 (August): 598–611. 10.1016/j.neuroimage.2017.06.040.

Hung, Lin W., Sophie Neuner, Jai S. Polepalli, et al. 2017. “Gating of Social Reward by Oxytocin in the Ventral Tegmental Area.” Science 357 (6358): 1406–11. 10.1126/science.aan4994.

Keysers, Christian, Ewelina Knapska, Marta A. Moita, and Valeria Gazzola. 2022. “Emotional Contagion and Prosocial Behavior in Rodents.” Trends in Cognitive Sciences 26 (8): 688–706. 10.1016/j.tics.2022.05.005.

Khani, Abbas, and Gregor Rainer. 2016. “Neural and Neurochemical Basis of Reinforcement-Guided Decision Making.” Journal of Neurophysiology 116 (2): 724–41. 10.1152/jn.01113.2015.

Lammel, Stephan, Byung Kook Lim, and Robert C. Malenka. 2014. “Reward and Aversion in a Heterogeneous Midbrain Dopamine System.” Neuropharmacology 76 (January): 351–59. 10.1016/j.neuropharm.2013.03.019.

Lee, E., I. Rhim, J. W. Lee, et al. 2016. “Enhanced Neuronal Activity in the Medial Prefrontal Cortex during Social Approach Behavior.” Journal of Neuroscience 36 (26): 6926–36. 10.1523/JNEUROSCI.0307-16.2016.

Li, S. William, Omer Zeliger, Leah Strahs, et al. 2022. “Frontal Neurons Driving Competitive Behaviour and Ecology of Social Groups.” Nature 603 (7902): 661–66. 10.1038/s41586-021-04000-5.

Misiołek, Klaudia, Marta Klimczak, Magdalena Chrószcz, et al. 2023. “Prosocial Behavior, Social Reward and Affective State Discrimination in Adult Male and Female Mice.” Scientific Reports 13 (1): 5583. 10.1038/s41598-023-32682-6.

Padilla-Coreano, Nancy, Kanha Batra, Makenzie Patarino, et al. 2022. “Cortical Ensembles Orchestrate Social Competition through Hypothalamic Outputs.” Nature 603 (7902): 667–71. 10.1038/s41586-022-04507-5.

Phillips, Mary L., Holly Anne Robinson, and Lucas Pozzo-Miller. 2019. “Ventral Hippocampal Projections to the Medial Prefrontal Cortex Regulate Social Memory.” eLife 8 (May): e44182. 10.7554/eLife.44182.

Pozo, Macarena, Maria Milà-Guasch, Roberta Haddad-Tóvolli, et al. 2023. “Negative Energy Balance Hinders Prosocial Helping Behavior.” Proceedings of the National Academy of Sciences 120 (15): e2218142120. 10.1073/pnas.2218142120.

Rault, Jean-Loup. 2019. “Be Kind to Others: Prosocial Behaviours and Their Implications for Animal Welfare.” Applied Animal Behaviour Science 210 (January): 113–23. 10.1016/j.applanim.2018.10.015.

Rogers-Carter, Morgan M., and John P. Christianson. 2019. “An Insular View of the Social Decision-Making Network.” Neuroscience & Biobehavioral Reviews 103 (August): 119–32. 10.1016/j.neubiorev.2019.06.005.

Rogers-Carter, Morgan M., Anthony Djerdjaj, K. Bates Gribbons, Juan A. Varela, and John P. Christianson. 2019. “Insular Cortex Projections to Nucleus Accumbens Core Mediate Social Approach to Stressed Juvenile Rats.” The Journal of Neuroscience 39 (44): 8717–29. 10.1523/JNEUROSCI.0316-19.2019.

Rogers-Carter, Morgan M., Juan A. Varela, Katherine B. Gribbons, et al. 2018. “Insular Cortex Mediates Approach and Avoidance Responses to Social Affective Stimuli.” Nature Neuroscience 21 (3): 404–14. 10.1038/s41593-018-0071-y.

Sapolsky, Robert M. 2005. “The Influence of Social Hierarchy on Primate Health.” Science 308 (5722): 648–52. 10.1126/science.1106477.

Scheggia, Diego, Filippo La Greca, Federica Maltese, et al. 2022. “Reciprocal Cortico-Amygdala Connections Regulate Prosocial and Selfish Choices in Mice.” Nature Neuroscience 25 (11): 1505–18. 10.1038/s41593-022-01179-2.

Shan, Qiang, You Hu, Shijie Chen, and Yao Tian. 2022. “Nucleus Accumbens Dichotomically Controls Social Dominance in Male Mice.” Neuropsychopharmacology 47 (3): 776–87. 10.1038/s41386-021-01220-1.

Smith, Monique L., Naoyuki Asada, and Robert C. Malenka. 2021. “Anterior Cingulate Inputs to Nucleus Accumbens Control the Social Transfer of Pain and Analgesia.” Science 371 (6525): 153–59. 10.1126/science.abe3040.

Soares-Cunha, Carina, Nivaldo A. P. De Vasconcelos, Bárbara Coimbra, et al. 2020. “Nucleus Accumbens Medium Spiny Neurons Subtypes Signal Both Reward and Aversion.” Molecular Psychiatry 25 (12): 3241–55. 10.1038/s41380-019-0484-3.

Song, Da, Chunjian Wang, Yue Jin, et al. 2023a. “Mediodorsal Thalamus-Projecting Anterior Cingulate Cortex Neurons Modulate Helping Behavior in Mice.” Current Biology 33 (20): 4330–4342.e5. 10.1016/j.cub.2023.08.070.

Van Loo, P. L. P., E. Van Der Meer, C. L. J. J. Kruitwagen, J. M. Koolhaas, L. F. M. Van Zutphen, and V. Baumans. 2003. “Strain-specific Aggressive Behavior of Male Mice Submitted to Different Husbandry Procedures.” Aggressive Behavior 29 (1): 69–80. 10.1002/ab.10035.

Wang, Fei, Jun Zhu, Hong Zhu, Qi Zhang, Zhanmin Lin, and Hailan Hu. 2011. “Bidirectional Control of Social Hierarchy by Synaptic Efficacy in Medial Prefrontal Cortex.” Science 334 (6056): 693–97. 10.1126/science.1209951.

Wu, Ye Emily, James Dang, Lyle Kingsbury, et al. 2021. “Neural Control of Affiliative Touch in Prosocial Interaction.” Nature 599 (7884): 262–67. 10.1038/s41586-021-03962-w.

Wu, Ye Emily, and Weizhe Hong. 2022. “Neural Basis of Prosocial Behavior.” Trends in Neurosciences 45 (10): 749–62. 10.1016/j.tins.2022.06.008.

Yamagishi, Atsuhito, Joungmin Lee, and Nobuya Sato. 2020. “Oxytocin in the Anterior Cingulate Cortex Is Involved in Helping Behaviour.” Behavioural Brain Research 393 (September): 112790. 10.1016/j.bbr.2020.112790.

Zalesky, Andrew, Alex Fornito, and Edward T. Bullmore. 2010. “Network-Based Statistic: Identifying Differences in Brain Networks.” NeuroImage 53 (4): 1197–207. 10.1016/j.neuroimage.2010.06.041.

Zhang, Mingmin, Ye Emily Wu, Mengping Jiang, and Weizhe Hong. 2024. “Cortical Regulation of Helping Behaviour towards Others in Pain.” Nature 626 (7997): 136–44. 10.1038/s41586-023-06973-x.

Zhou, Tingting, Carmen Sandi, and Hailan Hu. 2018. “Advances in Understanding Neural Mechanisms of Social Dominance.” Current Opinion in Neurobiology 49 (April): 99–107. 10.1016/j.conb.2018.01.006.

