## supplementary material for "Mouse strain and network-level activity differences underlie social decision-making"

Figure S1

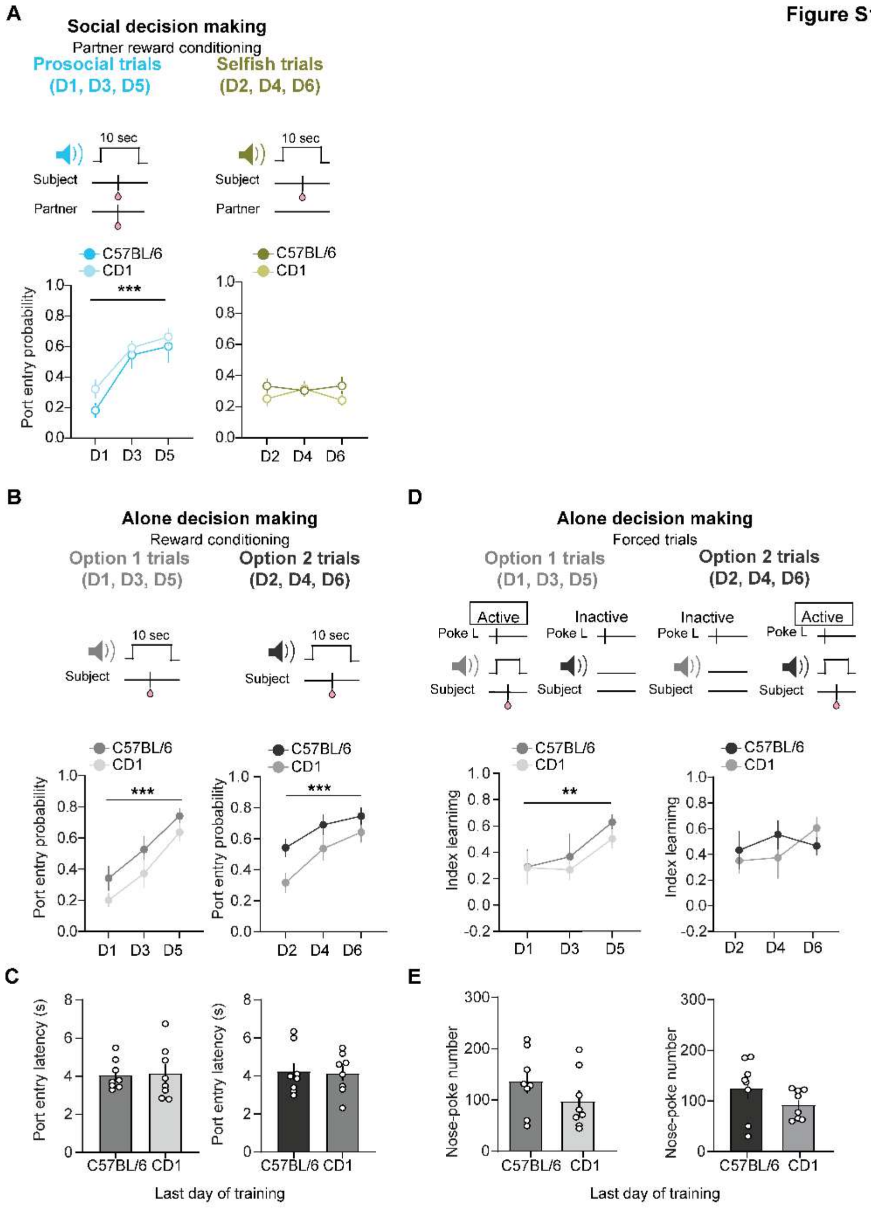


**Figure S1. Learning performance of partner mice and subjects trained alone.**
**A.** Top. Protocol training**.** Reward port entry probability during prosocial and selfish trials for C57BL/6 and CD1 partner mice under social decision-making (two-way RM ANOVA; Prosocial: strain, $F(1,14)= 1.48$, $p=0.24$; days, $F(1.18, 16.73)= 14.65$, $p= 0.0002$; interaction, $F (2, 28)= 0.29$, $p= 0.74$; Selfish: strain, $F(1,14)=1.81$, $p= 0.19$, days, $F(1.83, 25.64)= 0.13$, $p= 0.85$; interaction, $F(2, 28)= 0.92$, $p= 0.40$) **B.** Reward conditioning for C57BL/6 and CD1 mice trained alone. Top, protocol training. Bottom, reward port entry probability during CS-Option 1and CS-Option 2 trials showed no strain differences across days (two-way RM ANOVA; CS-Option 1 trials: strain, $F(1,14) = 3.41$, $p = 0.08$; days, $F(1.88, 26.35) = 24.59$, $p < 0.001$; interaction, $F(2,28) = 0.09$, $p = 0.91$; CS-Option 2 trials: strain, $F(1,14) = 3.93$, $p = 0.06$; time, $F(1.88, 26.42) = 27.25$, $p < 0.001$; interaction, $F(2,28) = 1.22$, $p = 0.30$). **C.** Reward port entry latencies during CS-Option 1

and CS-Option 2 for alone-trained mice do not show strain differences during the last day of training (unpaired t student test: prosocial, $t(14) = 0.1$, $p= 0.84$; selfish, $t(14) = 0.16$, $p= 0.87$). **D.** Forced-choice training during alone decision-making. Top, Protocol training. Bottom, Index learning indicates discrimination between active and inactive nose-pokes during CS-Option 1 and CS-Option 2 trials across strains of mice (Option 1: two-way RM ANOVA; strain, $F(1, 14) = 0.35$, $p = 0.56$; days, $F(2, 28) = 6.61$, $p = 0.004$; interaction, $F(2, 28) = 0.27$, $p = 0.76$; Option 2: two-way RM ANOVA; strain, $F(1, 14) = 0.14$, $p = 0.71$; days, $F(2, 28) = 0.86$, $p = 0.43$; interaction, $F(2, 28) = 1.12$, $p = 0.34$). **E.** Numbers of active vs inactive nose-poke responses on the last day of training for each trial type (unpaired t student test: option1, $t(14) = 1.22$, $p= 0.24$; option 2, $t(14) = 0.02$, $p= 0.98$).

Figure S2

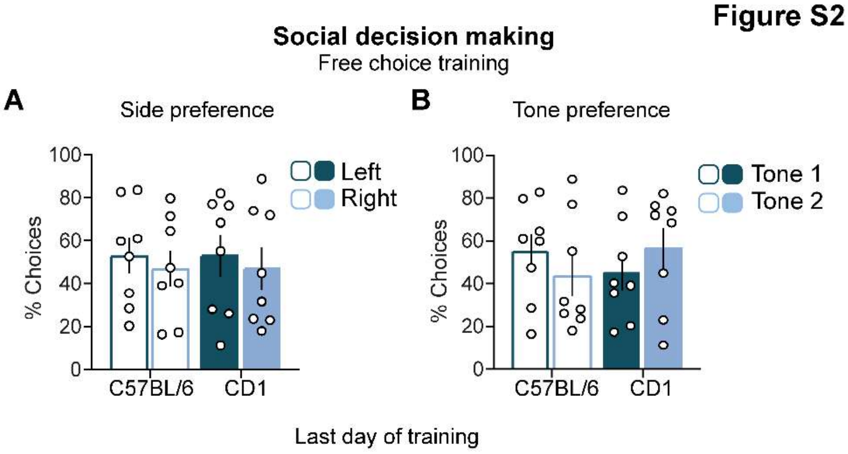


**Figure S2. No intrinsic side or tone preference during social decision-making.**
**A.** Side preference during social decision-making. On the final day of free-choice training, the percentage of choices did not differ between left and right nose-pokes across mouse strains. (two-way RM ANOVA; strain, $F(1, 14)= 0.00$, $p=0.99$; nose-poke side, $F(1,14)= 0.44$, $p= 0.51$; interaction, $F(1,14)= 0.00$, $p= 0.99$). **B,** Tone preference during social decision-making. On the final day of free-choice training, the percentage of choices did not differ between tones across mouse strains (two-way RM ANOVA; strain, $F(1, 14)= 0.09$, $p=0.75$; tones, $F(1,14)= 0.00$, $p= 0.99$; interaction, $F(1,14)= 0.97$, $p= 0.34$).

Figure S3

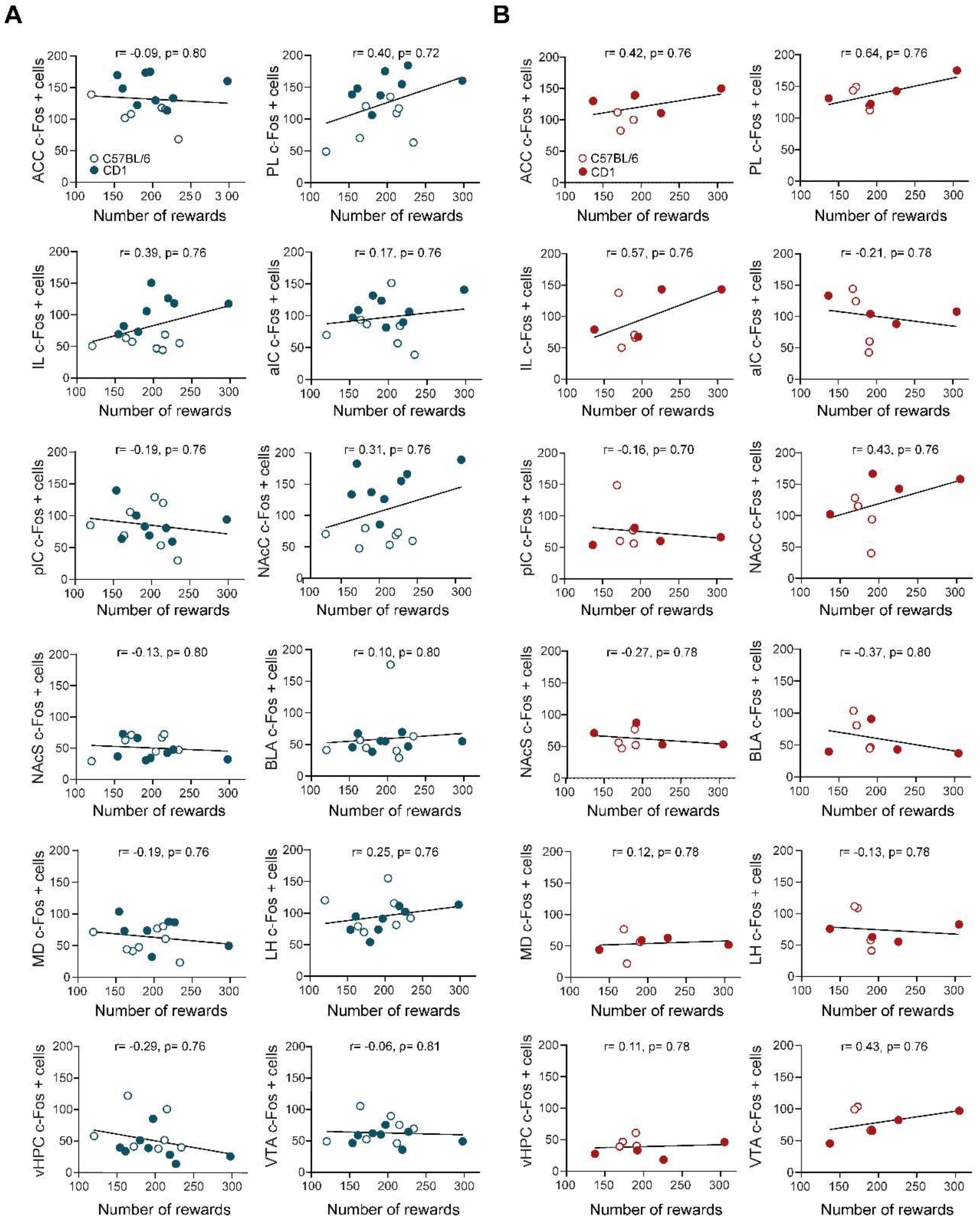


**Figure S3. Number of rewards obtained does not correlate with c-Fos activity.** **A**. Cortical and subcortical c-Fos–positive cell counts as a function of rewards consumed during social decision-making across all mice. **B.** Cortical and subcortical c-Fos–positive cell counts as a function of rewards consumed during alone decision-making across all mice.

Figure S4

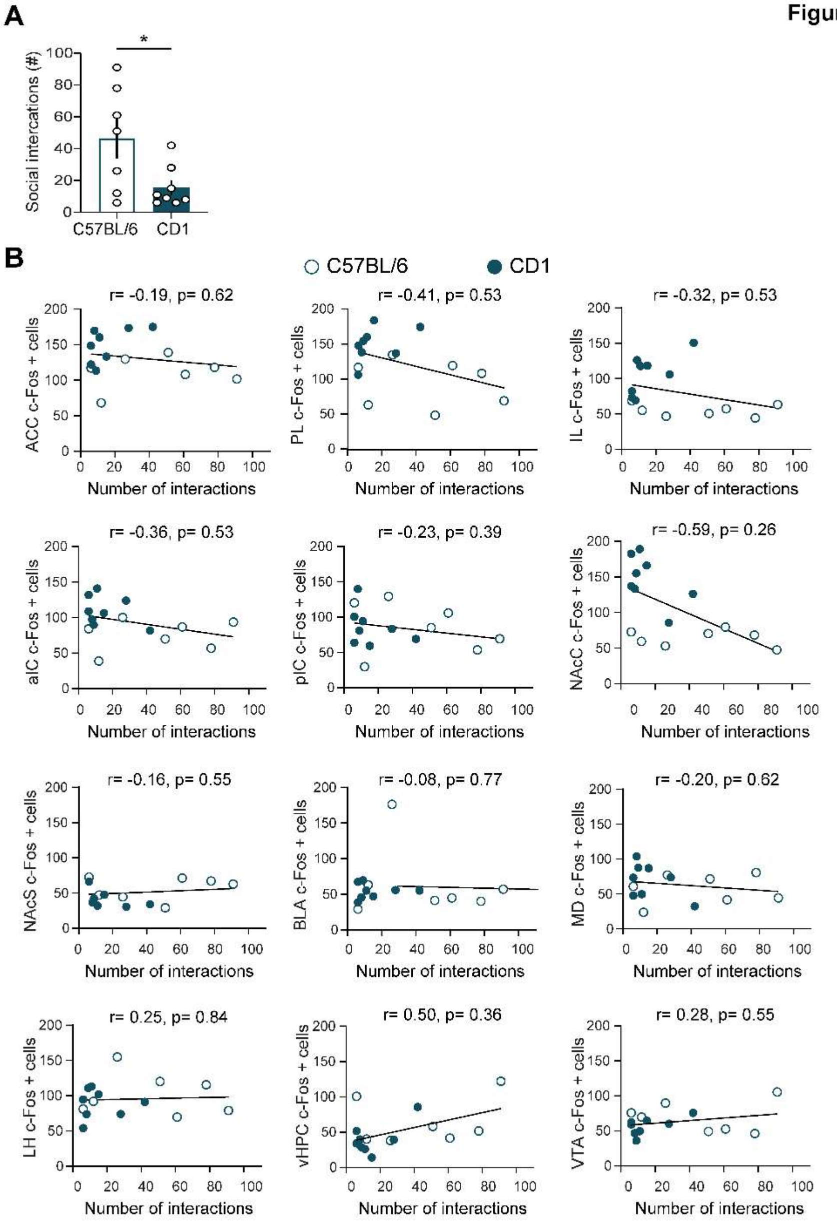


**Figure S4. Increased social interaction in C57BL/6 mice is not associated with c-Fos activity.**

**A.** Number of social interactions during the c-Fos test in C57BL/6 and Cd1 mice (unpaired t-test, $t(13)= 2.46$, $p=0.02$). **B.** Cortical and subcortical c-Fos+ cells count as a function of the number of social interactions during social decision-making across all mice.

Figure S5

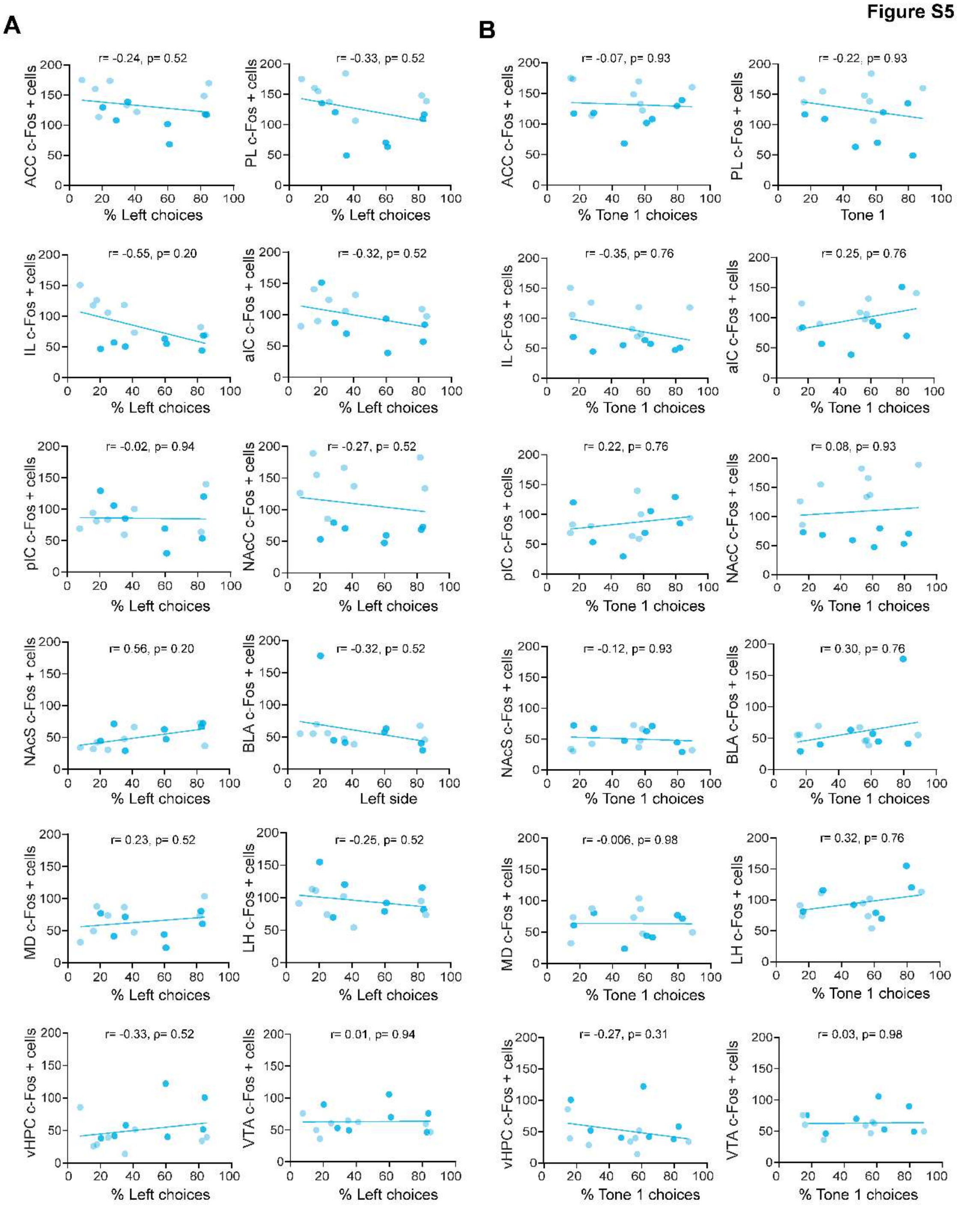


**Figure S4. Side- and tone-guided choices do not correlate with c-Fos activity during social decision-making.**

**A**. Cortical and subcortical c-Fos–positive cell counts as a function of the percentage of left nose-poke choices during social conditions across all mice. **B.** Cortical and subcortical c-Fos–positive cell counts as a function of the percentage of tone 1 choices during social decision-making across all mice.

Figure S6

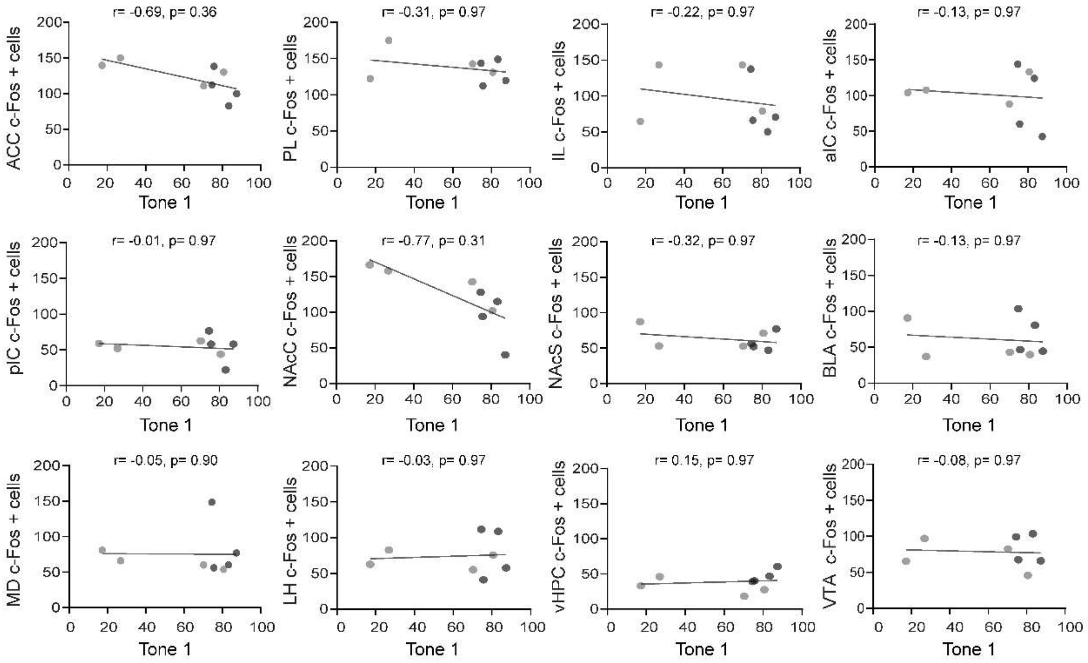


**Figure S6. Tone-guided choices do not correlate with c-Fos activity in the alone condition.**

Cortical and subcortical c-Fos–positive cell counts as a function of the percentage of tone 1 choices during alone decision-making across all mice.

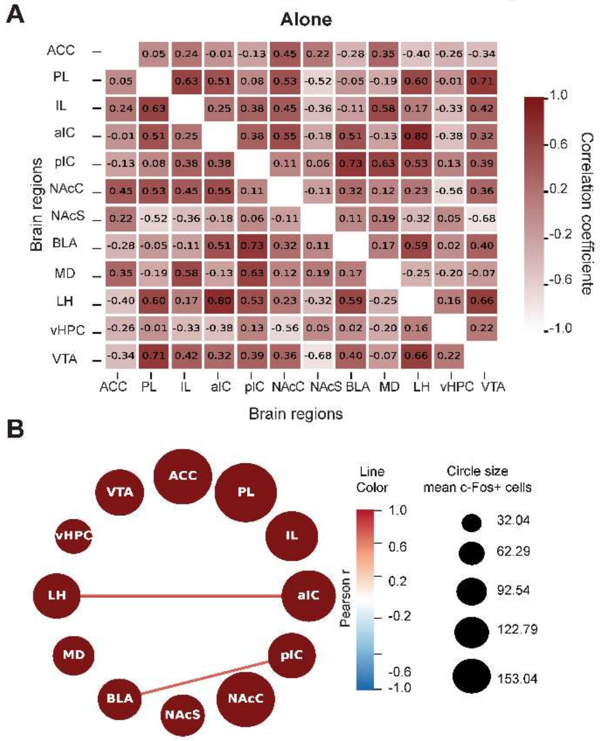


**Figure S7. Cross-regional c-Fos correlations during alone decision-making.**
**A.** Heatmap showing pairwise Pearson correlations of c-Fos+ cell counts across brain regions during the alone condition. **B**. Network representation illustrating suprathreshold cross-regional associations during alone decision-making for all mice. Network visualizations show significant cross-correlations for each condition based on a null-distribution method. Line color represents Pearson r value. Only cross-correlations in the top/bottom 2.5% of null distribution are shown (n= 4 C57 and 4 CD1).
